# Pulmonary colonization of *Candida albicans* drives breast cancer lung metastasis via candidalysin

**DOI:** 10.64898/2026.10.23.759946

**Authors:** Zhongyi Ma, Lei Chen, Yike Fang, Feiyan Gao, Lihan Liu, Jin Wu, Hongdan Chen, Yueru Tian, Yuping Zhang, Xinlin Yang, Yumei Wang, Mengdi Ren, Yulan Liu, Lin Zhu, Changbin Chen, Ming Guan, Peng Tang, Linqi Wang, Chong Li

## Abstract

The intratumoral mycobiome is emerging as a key modulator of cancer progression, with specific fungal taxa as potential biomarkers. However, whether and how mycobiomes colonizing distant organs contribute to tumor evolution remains largely unexplored. Here, we identify selective enrichment of *Candida albicans* in breast cancer lung metastases. Mechanistically, the fungal virulence factor candidalysin activates pulmonary mast cells, which release chemotactic mediators that expand immunosuppressive populations and suppress cytotoxic lymphocyte activity, remodeling the lung into a metastasis-permissive microenvironment. Analysis of clinical isolates and isogenic mutants further demonstrates that candidalysin secretion levels dictate metastasis-promoting potential. Informed by these findings, we developed a combination regimen that concurrently eliminates fungal colonization, stabilizes mast cells, and reverses the immunosuppressive microenvironment, thereby significantly curtailing metastatic progression. Collectively, our study establishes pulmonary *C. albicans* colonization as a driver of breast cancer lung metastasis and provides a mechanistic framework for targeting mycobiome-host interactions in metastatic disease.

## Introduction

Breast cancer is the most frequently diagnosed malignancy among women worldwide, with an estimated 2.3 million new cases and approximately 670,000 related deaths annually, and the mortality burden remains largely unabated.^1–3^ The vast majority of breast cancer-related mortality is attributable to metastasis,^4–6^ an intricate process shaped by multiple regulatory factors, including tumor cell-inherent biological properties and the heterogeneous features of organ-specific niches.^7–10^ Seminal studies over the past several years have largely emphasized cancer cell-autonomous traits and their direct roles in driving metastatic progression.^11,12^ Cancer cell-secreted proteases can drive organ-specific metastasis by remodeling local and distal microenvironments.^13–15^ Additionally, cancer cells can deploy extracellular vesicles as vehicles for intercellular and interorgan communication, thereby establishing permissive pre-metastatic niches in distant organs and promoting metastatic relapse.^16–19^ Nevertheless, beyond the tumor cell-intrinsic mechanisms that influence the pre-metastatic microenvironment, the role of organ-specific niches in metastatic colonization and outgrowth remains largely unexplored.

The intratumoral microbiome has emerged as a key component of the microenvironment, profoundly shaping tumor initiation, progression, metastasis, and therapeutic response.^20–23^ Elucidating how the microbiome interacts with tumor immunity has therefore become a major focus bridging basic and translational cancer research. As a minor yet highly immunoreactive component of the human microbiome, commensal fungi critically regulate tumor homeostasis.^24–26^ Recent studies have begun to chart tumor-associated mycobiota and have revealed mechanistic links between specific fungi and cancer progression.^24,26–30^ Specifically, these fungi often exert pro-tumorigenic effects by reprogramming the local immune microenvironment. In pancreatic cancer, intratumor *Malassezia* promotes tumorigenesis by activating the C3 complement cascade and inducing type 2 immunity,^26^ whereas the tumor-resident fungus *Aspergillus sydowii* accelerates lung cancer progression by expanding myeloid-derived suppressor cells.^30^ While these findings highlight the significant local impact of tumor-resident fungi, whether commensal fungi colonizing distal organs contribute to metastatic progression has not been systematically investigated.

Here, focusing on the lung, the predominant site of breast cancer dissemination and a mucosal organ harboring a resident mycobiome, we identify selective enrichment of *C. albicans* within lung metastatic lesions of breast cancer patients. Mechanistically, *C. albicans* secretes the virulence factor candidalysin to activate lung mast cells, triggering their degranulation and chemokine release that promotes the expansion of immunosuppressive populations and suppresses cytotoxic lymphocyte function. This cascade establishes an immunosuppressive pulmonary niche that directly supports metastatic colonization and outgrowth. Notably, clinical *C. albicans* isolates exhibit marked heterogeneity in candidalysin secretion, and the secretion capacity of this virulence factor positively correlates with metastasis-promoting potential. These findings prompted us to develop a combination therapy that targets this candidalysin-driven mast cell cascade and the resulting immunosuppressive niche, which conferred potent protection against breast cancer lung metastasis in preclinical models. Together, our study uncovers a mechanism by which lung-colonizing fungi drive breast cancer metastasis and provides a rationale for patient risk stratification and targeted therapeutic interventions.

## Results

### Breast cancer lung metastases exhibit fungal dysbiosis and enrichment of *C. albicans*

To characterize the pulmonary mycobiome in the context of breast cancer metastasis, we performed ITS sequencing on 157 lung specimens from multiple clinical centers, including 112 histopathologically confirmed metastatic lesions and 45 non-metastatic control tissues (Figure 1A; Table S1). Given the inherent challenges of low-biomass samples,^30–32^ we implemented a rigorous quality-control pipeline incorporating environmental and reagent blank controls (Figure S1A). To accommodate the pronounced length polymorphism of ITS amplicons, DADA2 processing parameters were specifically optimized to enable accurate sequence denoising, paired-end read merging, and chimera removal. Stringent prevalence- and abundance-based filtering thresholds were subsequently applied to ensure high-confidence discrimination of endogenous fungal signals.

**Figure 1.**
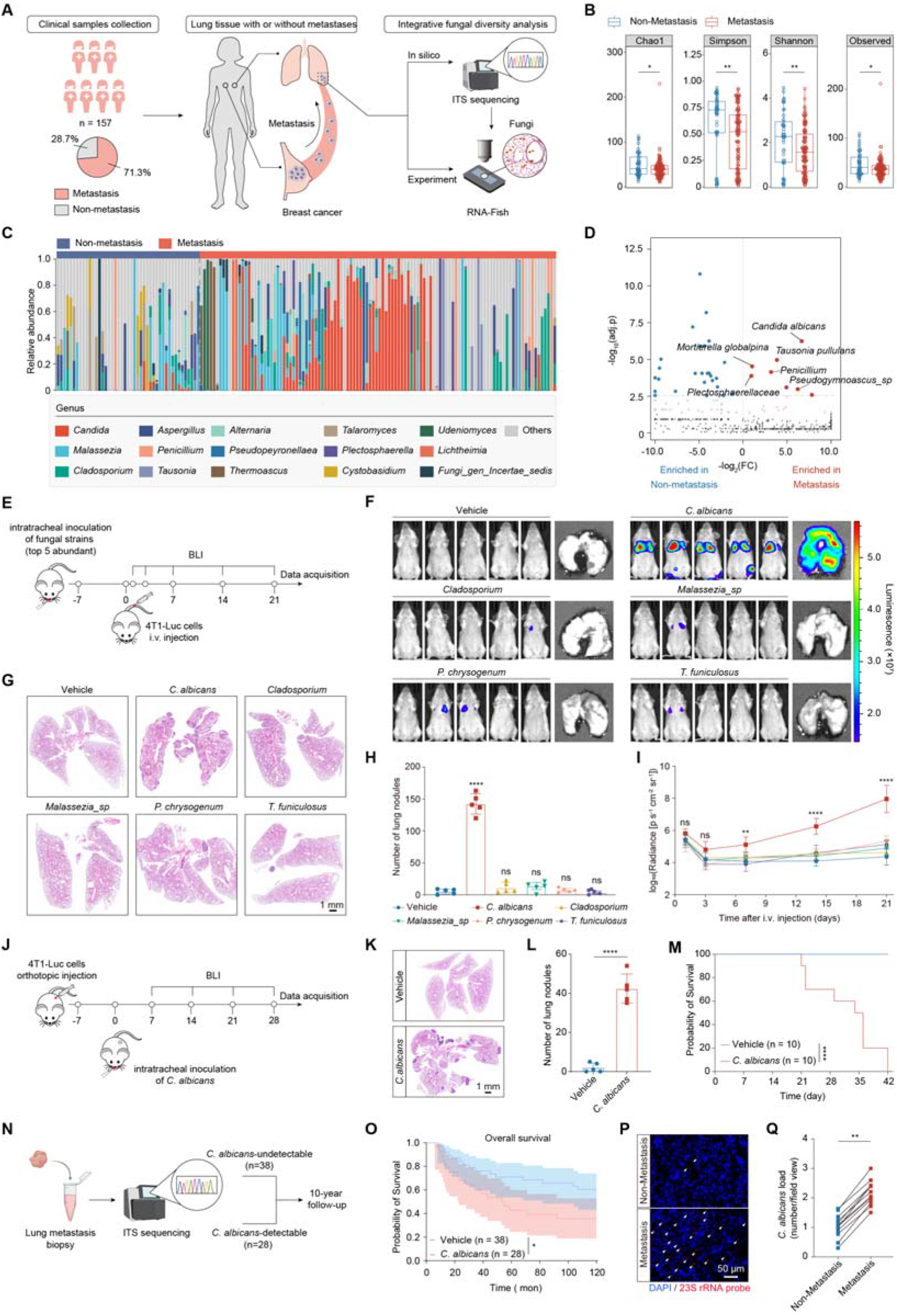
Pulmonary fungal dysbiosis and *C. albicans* enrichment promote breast cancer lung metastasis. (A) Schematic overview of clinical sample collection and integrated fungal diversity analysis. A total of 157 multicenter clinical pulmonary biospecimens were included in this study. Baseline characteristics of enrolled participants are summarized in Table S1. (B) Comparison of pulmonary fungal alpha diversity between the non-metastasis and metastasis groups. (C) Genus-level distribution of pulmonary fungal community structures in the non-metastasis and metastasis groups. (D) Volcano plot illustrating the differentially abundant fungal taxa between the metastasis and non-metastasis groups. (E) Schematic of the animal experimental workflow used to evaluate the effects of pulmonary colonization by distinct fungal species on breast cancer lung metastasis. (F–I) Evaluation of the metastasis-promoting effects of the five most abundant fungal strains. (F) Representative BLI images of pulmonary metastases. Images of mice at day 21 and representative *ex vivo* lungs from each group are shown. Complete longitudinal tracking series across all time points and full *ex vivo* lung panels are provided in Figure S2A. (G) Representative H&E-stained images. Scale bar, 1 mm. (H) quantification of pulmonary metastatic nodules. (I) Longitudinal bioluminescence tracking of mice colonized by the respective fungal strains. (J) Schematic of the animal experimental workflow used to evaluate the effect of pulmonary *C. albicans* colonization on spontaneous breast cancer lung metastasis. (K–L) Representative H&E-stained images (K) and quantification of pulmonary metastatic nodules (L). (M) Kaplan-Meier survival curves of mice in the vehicle and *C. albicans* groups (n = 10). (N–O) Evaluation of patient survival outcomes based on *C. albicans* detectability in lung metastasis samples. (N) Schematic of the clinical cohort analysis. Lung metastasis samples underwent ITS sequencing, stratifying patients into *C. albicans*-undetectable (n = 38) and *C. albicans*-detectable (n = 28) groups for a 10-year follow-up. (O) Kaplan-Meier survival curves. (P–Q) Representative FISH images (P) and quantification (Q) of *C. albicans* in lung metastatic lesions and paired non-metastatic tissues. Data are presented as mean ± SEM. ns, not significant; *p < 0.05; **p < 0.01; ****p < 0.0001 compared using unpaired Student’s t test (two-tailed) (B, L, and Q), one-way ANOVA (H and I), and log rank (Mantel-Cox) test (M, and O).

Metastatic lesions exhibited significantly reduced fungal α-diversity, with substantial reductions in Chao1, observed ASVs, Shannon, and Simpson indices compared with non-metastatic controls (Figure 1B; Figure S1B), indicating pronounced simplification and dysbiosis of the pulmonary fungal community. Jaccard distance-based beta-diversity analysis further revealed a clear separation in fungal community composition between metastatic and non-metastatic tissues (Figures S1C-S1D).

Moreover, within-group pairwise distances were significantly reduced among metastatic lesions (Figure S1E), indicating a convergent restructuring of the local fungal ecosystem during metastatic progression. Notably, *Candida* exhibited pronounced ecological dominance at the genus level in metastatic samples, substantially exceeding its relative abundance in non-metastatic controls (Figure 1C). Differential abundance analysis further identified *C. albicans* as the most significantly enriched fungal species in metastatic lesions, surpassing all other differentially represented taxa (Figure 1D; Figure S1F). These data uncover a pathological simplification of the pulmonary fungal landscape and define a distinctive microbial signature of the breast cancer lung metastatic microenvironment.

### Lung colonization by *C. albicans* enhances metastatic outgrowth

To systematically validate the five most abundant fungal species *in vivo*, we intratracheally inoculated mice with each species and then challenged them intravenously with 4T1-Luc breast cancer cells (Figure 1E). Among the species tested, only *C. albicans* colonization induced a robust pro-metastatic phenotype, increasing pulmonary bioluminescence by exceeding three orders of magnitude and elevating metastatic nodule counts by nearly 27-fold relative to controls (Figures 1F-1I; Figure S2A).

To further assess whether pulmonary *C. albicans* colonization promotes spontaneous lung metastasis, we first optimized the fungal inoculation dose to establish sustained pulmonary colonization without compromising baseline survival (Figures S2B-S2F). Mice bearing orthotopic 4T1-Luc mammary tumors were then intratracheally inoculated with the optimized dose of *C. albicans* at an early stage of tumor development (Figure 1J). In this model, *C. albicans* colonization minimally affected primary tumor growth (Figures S2G-2K) but profoundly promoted lung metastasis, as evidenced by a substantial increase in pulmonary bioluminescence (Figures S2L-S2M), extensive surface metastatic nodules (Figures 1K-1L), and markedly shortened survival (Figure 1M). *C. albicans* consequently functions as a pathogenic driver, rather than a passive bystander, of breast cancer metastatic progression.

### Pulmonary *C. albicans* burden associates with poor clinical outcome and lesion-specific enrichment

We explored the clinical relevance of these findings in a 10-year retrospective cohort of 66 patients with breast cancer lung metastasis to assess the association between pulmonary fungal burden and patient outcomes (Figure 1N; Table S2). Survival analysis revealed that pulmonary *C. albicans* abundance significantly stratified overall survival, providing clinical evidence linking fungal burden with cancer progression (Figure 1O). To spatially map fungal distribution in clinical specimens, fluorescence in situ hybridization (FISH) was performed, revealing pronounced spatial heterogeneity, with fungal burden significantly higher in metastatic lesions than in matched non-tumor tissues from the same patients (Figures 1P-1Q), consistent with preferential enrichment of *C. albicans* within metastatic niches. The concordance between systemic prognostic stratification and local spatial enrichment suggests that *C. albicans* is a functionally active microenvironmental component that is selectively enriched within metastatic lesions and contributes to disease progression.

### *C. albicans* colonization remodels the pulmonary immunosuppressive landscape and functionally reprograms mast cells

While commensal microbes are increasingly recognized to drive cancer progression through host immune remodeling,^33,34^ whether pulmonary *C. albicans* colonization promotes breast cancer metastasis through such immune modulation remains to be determined. To address this, we performed single-cell RNA sequencing (scRNA-seq) on CD45 immune cells sorted from non-metastatic lungs of orthotopic 4T1 tumor-bearing mice with established pulmonary *C. albicans* colonization.

Temporal analysis showed that no overt metastatic lesions were detectable in the distal lungs within 14 days after orthotopic tumor implantation (Figure S3A), thereby defining day 14 or earlier as the pre-metastatic window in the 4T1 model. Importantly, single-cell profiling revealed that the pulmonary immune landscape had already undergone profound remodeling within this pre-metastatic interval (Figures 2A-2B; Figures S3B-S3G). Ro/e analyses revealed a marked remodeling of the pulmonary immune microenvironment following *C. albicans* colonization. Specifically, neutrophils and PMN-MDSCs underwent a substantial expansion, whereas pulmonary infiltration of CD4 and CD8 T cells, NK cells, and dendritic cells (DCs) was markedly reduced (Figures 2B-2C).

**Figure 2.**
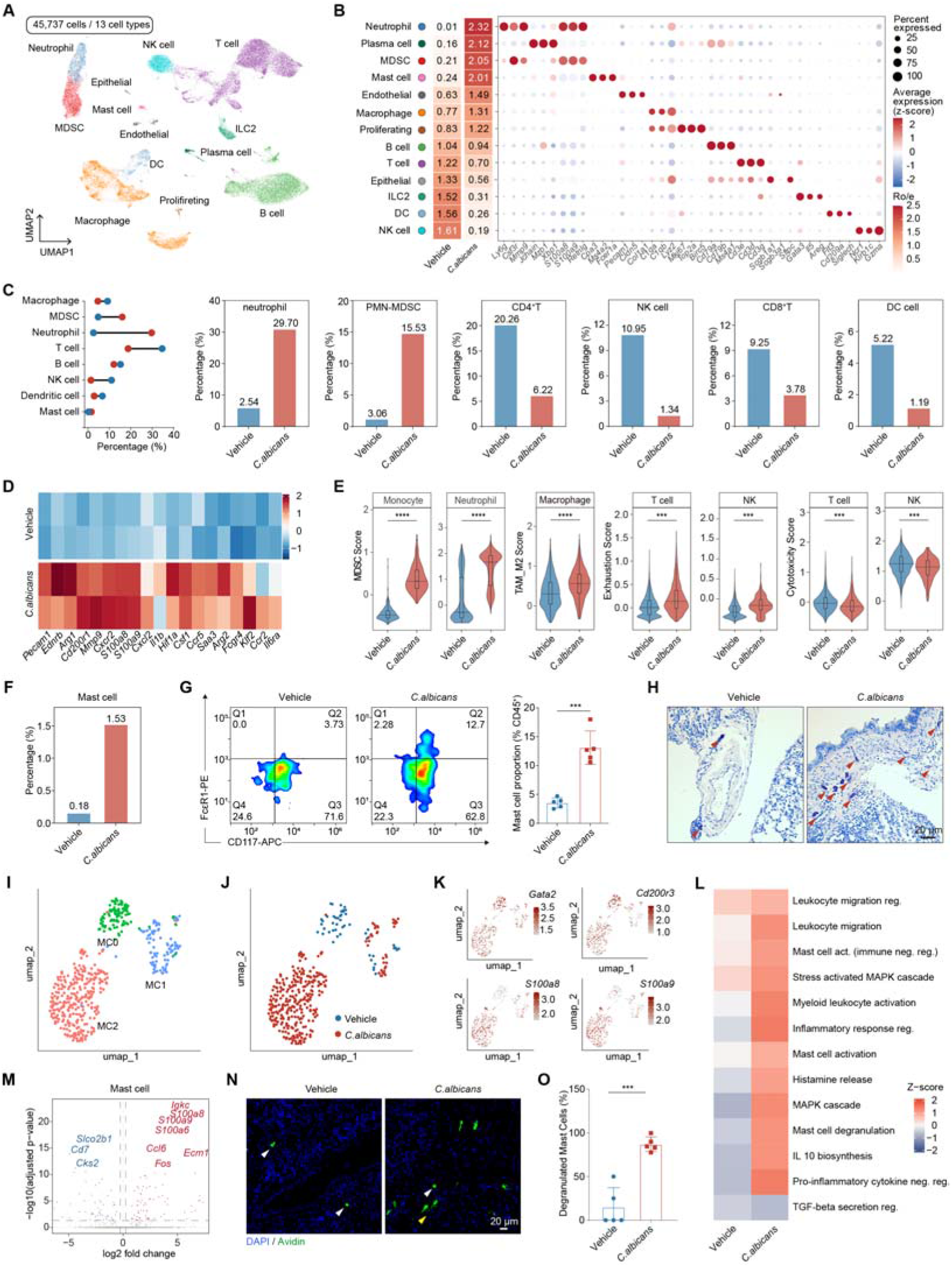
Immunosuppressive remodeling and mast cell activation upon pulmonary *C. albicans* colonization. (A) UMAP visualization of scRNA-seq profiles showing major cell clusters. (B) Heatmap showing the distribution preferences of major pulmonary cell subsets across experimental groups (left) and dot plot showing the expression of representative marker genes (right). (C) Relative proportions of major immune cell subsets in mouse lungs. (D) Z-score heatmap of differentially expressed genes (DEGs) in pulmonary myeloid cells. (E) Violin plots showing feature scores in pulmonary myeloid and lymphoid cell subsets. (F) scRNA-seq-based quantification of changes in the proportion of pulmonary mast cells. (G–H) Flow cytometric analysis of pulmonary mast cells (G) and representative toluidine blue staining of lung tissues (H). (I) UMAP visualization showing the segregation of mast cells into three distinct subclusters (MC0, MC1, and MC2). (J) UMAP visualization as in (I), colored by experimental group to show the distribution of cells across mast cell subclusters. (K) UMAP feature plots showing the expression of selected marker genes within mast cell clusters. Color intensity indicates relative gene expression levels. (L) Heatmap showing pathway enrichment in mast cells from the Vehicle and *C. albicans* groups. (M) Volcano plot showing differentially expressed genes in mast cells. (N–O) Representative immunofluorescence images (N) and quantification (O) of degranulated mast cells in lung tissues. DAPI (blue) was used to counterstain nuclei, and Avidin (green) labeled mast cells. Yellow arrows indicate degranulated mast cells; white arrows indicate non-degranulated mast cells. Scale bar, 20 μm. Data are presented as mean ± SEM. ***p < 0.001; ****p < 0.0001 compared using unpaired Student’s *t* test (two-tailed) (E, G, and O).

To define the phenotypic and functional states of the recruited myeloid populations, their transcriptional programs and functional gene modules were further interrogated. Differential expression analysis revealed that *C. albicans* colonization induced a coordinated set of effector genes associated with immunosuppression and angiogenesis (*Arg1*, *Arg2*, *Pecam1*, and *Ednrb*),^35–37^ extracellular matrix remodeling (*Mmp9*),^38^ inflammatory and chemotactic signaling (*Cxcr2*, *S100a8*, *S100a9*, *Cxcl2*, and *Il1b*),^39–41^ and hypoxic adaptation (*Hif1a*) (Figure 2D). Module scoring and pathway analyses revealed that *C. albicans* colonization markedly increased MDSC signature scores in monocytes and neutrophils and induced an M2-like macrophage signature in macrophages (Figure 2E). Concomitantly, T cells and NK cells exhibited diminished cytotoxicity signatures and enhanced exhaustion-associated programs, indicative of profound lymphoid dysfunction (Figure 2E).

Within this dynamically remodeled immune landscape, mast cells emerged as a previously overlooked immune population in conventional immunological studies.^42–44^ Single-cell profiling defined this population by selective expression of canonical mast cell lineage genes, including *Cpa3*, *Kit*, *Mcpt8*, *Ms4a2*, *Fcer1a*, and *Gata2* (Figure S3H),^45^ and revealed a marked expansion of mast cells following *C. albicans* colonization (Figure 2F). Flow cytometry and in situ histological staining further confirmed a substantial increase in pulmonary mast cell abundance in colonized mice, with prominent accumulation within perivascular and peribronchial interstitial regions (Figures 2G-2H).

To further resolve the functional heterogeneity of mast cells during remodeling of the pro-metastatic microenvironment, mast cells were subjected to high-resolution unsupervised subclustering, which identified three transcriptionally distinct states (MC0, MC1, and MC2) (Figure 2I; Figure S3I).^46^ Cell-origin mapping revealed a pronounced phenotypic shift following *C. albicans* colonization. Whereas mast cells in vehicle-treated lungs were predominantly composed of MC0 (*Gata2, Slc18a2, Cd200r3*) and the mature MC1 (*Mcpt1, Mcpt2, Cma1, Tpsb2, Tpsab1, Tph1*) population, pulmonary *C. albicans* colonization disrupted this steady-state distribution and selectively drove polarization and extensive expansion of the inflammatory/microenvironment-remodeling MC2 (*S100a8, S100a9, Ccl6, Fos, Anxa1, Dusp1*) subset, which became the dominant mast cell population in colonized lungs (Figures 2J-2K).

GO enrichment analysis further showed that genes upregulated in mast cells from colonized lungs were significantly enriched in MAPK signaling, leukocyte migration and recruitment, mast cell degranulation, and IL-10 biosynthetic processes (Figure 2L). The inflammatory and pro-metastatic mediators *S100a8*, *S100a9*, and *S100a6*, together with the chemokine *Ccl6*, were markedly upregulated following colonization (Figure 2M). Furthermore, avidin fluorescence staining revealed intense avidin-positive signals accompanied by extracellular granule deposition in infiltrating mast cells,^47,48^ providing in situ evidence that pulmonary *C. albicans* colonization induces robust mast cell activation and degranulation (Figures 2N-2O).

Potentially reflecting the functional consequences of this robust activation, CellChat analysis revealed extensive mast cell–myeloid communication, with overall interaction strength markedly increased relative to controls. Mast cells also established prominent signaling with lymphoid populations through the IL-4–IL-4R axis and engaged Tregs through the CD274–PDCD1 and CD86–CTLA4 axes, consistent with enhanced negative immune regulatory signaling (Figure 2M; Figures S3J-S3K). Collectively, these results indicate that distal *C. albicans* colonization reshapes the pre-metastatic pulmonary immune landscape through coordinated reprogramming of suppressive myeloid cells, exacerbation of lymphocyte exhaustion, and expansion and polarization of mast cells, thereby establishing an immune environment permissive for subsequent breast cancer lung metastasis.

### Mast cells are required for *C. albicans*-driven pulmonary immunosuppression and metastatic progression

To define how *C. albicans* reshapes the pulmonary immune microenvironment, we focused on mast cells,^49,50^ which exhibited pronounced expansion and phenotypic polarization in the preceding analyses. *C. albicans* challenge promoted mast cell migration and degranulation (Figures S4A-S4B), and markedly increased the secretion of pro-inflammatory and chemotactic cytokines, including CCL2, CCL3, CXCL2, IL-1β, IL-6, and TNF-α (Figure 3A).^51,52^

**Figure 3.**
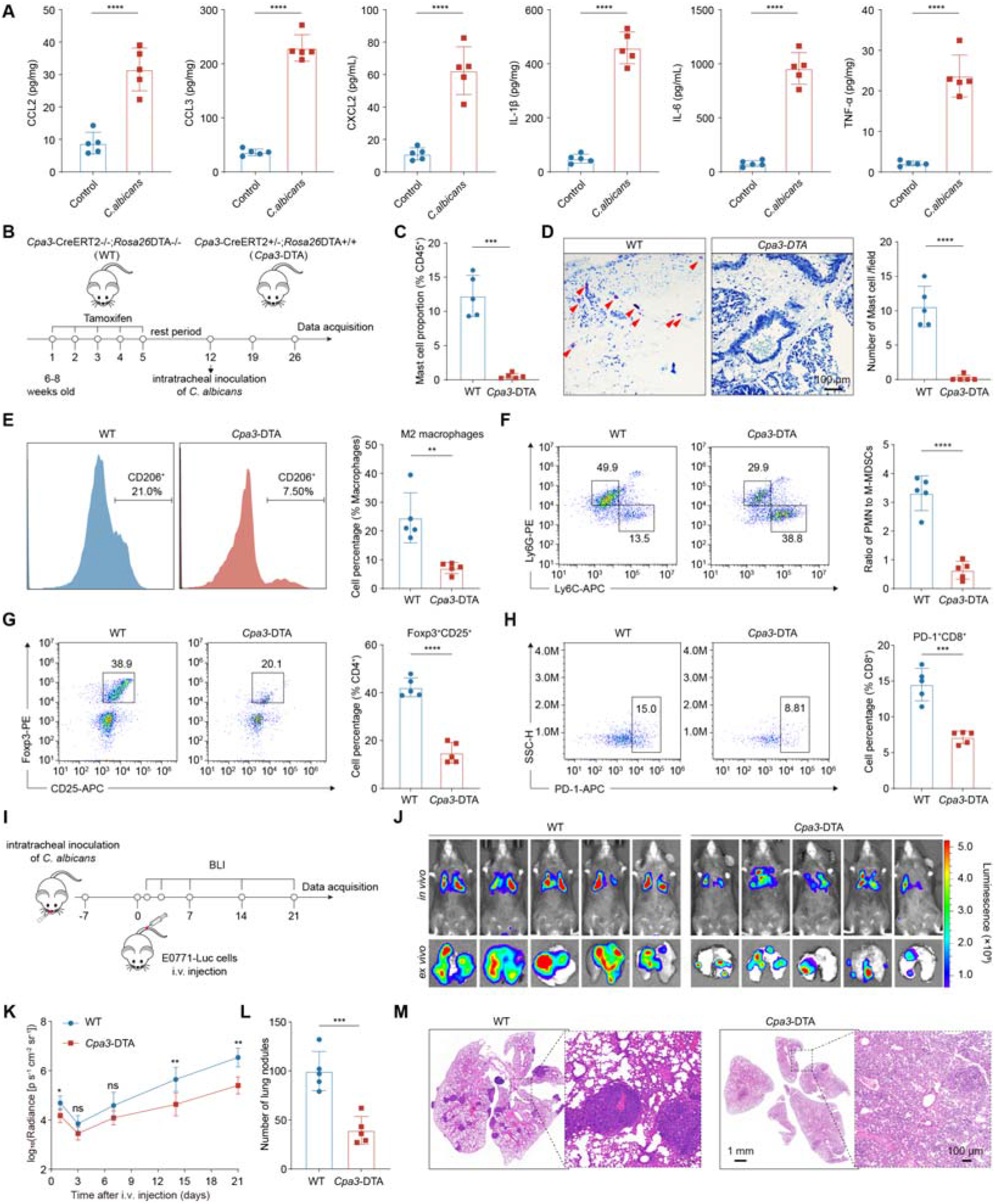
Mast cells are required for C. albicans-driven pulmonary immunosuppression and breast cancer metastasis. (A) Cytokine levels in mast cell culture supernatants measured by ELISA. (B) Schematic illustrating the experimental workflow for conditional mast cell depletion using *Cpa3*-DTA mice followed by *C. albicans* inoculation. (C) Flow cytometric quantification of mast cell proportions in the lungs of WT and *Cpa3*-DTA mice. (D) Representative toluidine blue staining (left) and quantification of mast cells per field (right) in lung tissues. Scale bar, 100 μm. (E–H) Flow cytometric analysis of immune cell populations in lung tissues. Representative flow cytometry plots and statistical quantification of M2 macrophages (E), the ratio of PMN-MDSCs to M-MDSCs (F), Tregs (G), and PD-1^+^ CD8^+^ T cells (H). (I) Schematic of the experimental workflow for evaluating the effect of pulmonary *C. albicans* colonization on lung metastasis in WT and *Cpa3*-DTA mice. (J–K) Representative BLI images of pulmonary metastases (J) and longitudinal bioluminescence tracking of mice (K). (L–M) Quantification of pulmonary metastatic nodules (N) and representative H&E-stained images (O). Data are presented as mean ± SEM. ns, not significant; *p < 0.05; **p < 0.01; ***p < 0.001; ****p < 0.0001 compared using unpaired Student’s *t* test (two-tailed) (A, C-H, and K-L).

To establish the functional requirement for mast cells *in vivo*, a tamoxifen-inducible *Cpa3*-DTA conditional mast cell ablation model driven by the mast cell-associated carboxypeptidase A3 (*Cpa3*) locus was employed (Figures S4C-S4D). ^53,54^ After tamoxifen-induced depletion, WT and *Cpa3*-DTA mice were intratracheally inoculated with *C. albicans* to determine whether mast cell loss altered fungal colonization-induced remodeling of the pulmonary immune microenvironment (Figure 3B). Flow cytometry combined with toluidine blue histochemistry verified efficient mast cell depletion in *Cpa3*-DTA mice (Figures 3C-3D; Figure S4E). While *C. albicans* colonization drove pronounced mast cell expansion and tissue infiltration in wild-type lungs, this response was abolished in *Cpa3*-DTA mutants.

To assess how mast cell depletion affected the broader pulmonary immune landscape, major immune cell subsets were further profiled. The immunosuppressive network induced by pulmonary *C. albicans* colonization was markedly attenuated following mast cell ablation. Compared with WT controls, *Cpa3*-DTA mice exhibited substantially reduced proportions of M2-like macrophages, MDSCs, Tregs, and PD-1 CD8 T cells in the lung (Figures 3E-3H; Figures S4F-S4I). These findings indicate that mast cells are required for the establishment and maintenance of the immunosuppressive pulmonary microenvironment induced by fungal colonization.

To determine whether the metastasis-promoting effect of *C. albicans* was specifically dependent on mast cells, an experimental E0771-Luc lung metastasis model was subsequently established in WT and *Cpa3*-DTA mice with pre-established pulmonary *C. albicans* colonization (Figure 3I). Longitudinal bioluminescence imaging (BLI) revealed that selective mast cell ablation markedly restrained *C. albicans*-driven metastatic progression, with substantially reduced pulmonary bioluminescence signals in *Cpa3*-DTA mice compared with WT controls (Figures 3J-3K). Endpoint quantification of gross pulmonary metastatic nodules and histopathological assessment further confirmed that mast cell depletion substantially reduced the metastatic burden induced by fungal colonization (Figures 3L-3M). Collectively, these findings establish mast cell activation as a critical upstream event through which pulmonary *C. albicans* colonization orchestrates an immunosuppressive lung microenvironment and promotes breast cancer lung metastasis.

### Candidalysin is the key fungal effector that triggers mast cell activation

To identify the specific fungal component that triggers mast cell activation during *C. albicans* colonization, we devised a stepwise screening strategy (Figure 4A). We first assessed whether degranulation required live fungi, direct cell–cell contact, or soluble proteinaceous factors. Toluidine blue staining and β-hexosaminidase (β-Hex) release assays showed that cell-free fungal culture supernatant was sufficient to induce robust mast cell degranulation (Figure 4B; Figure S5A). Ultrafiltration localized the stimulatory activity predominantly to the 2-5 kDa fraction (Figure 4B; Figure S5A). Peptidomic analysis coupled with secretome database cross-referencing identified candidalysin as the most abundant polypeptide within the active fraction, with a molecular mass of 3.31kDa, nominating this *ECE1*-encoded cytolytic peptide as a candidate mediator of mast cell activation (Figure 4C). ^55,56^

**Figure 4.**
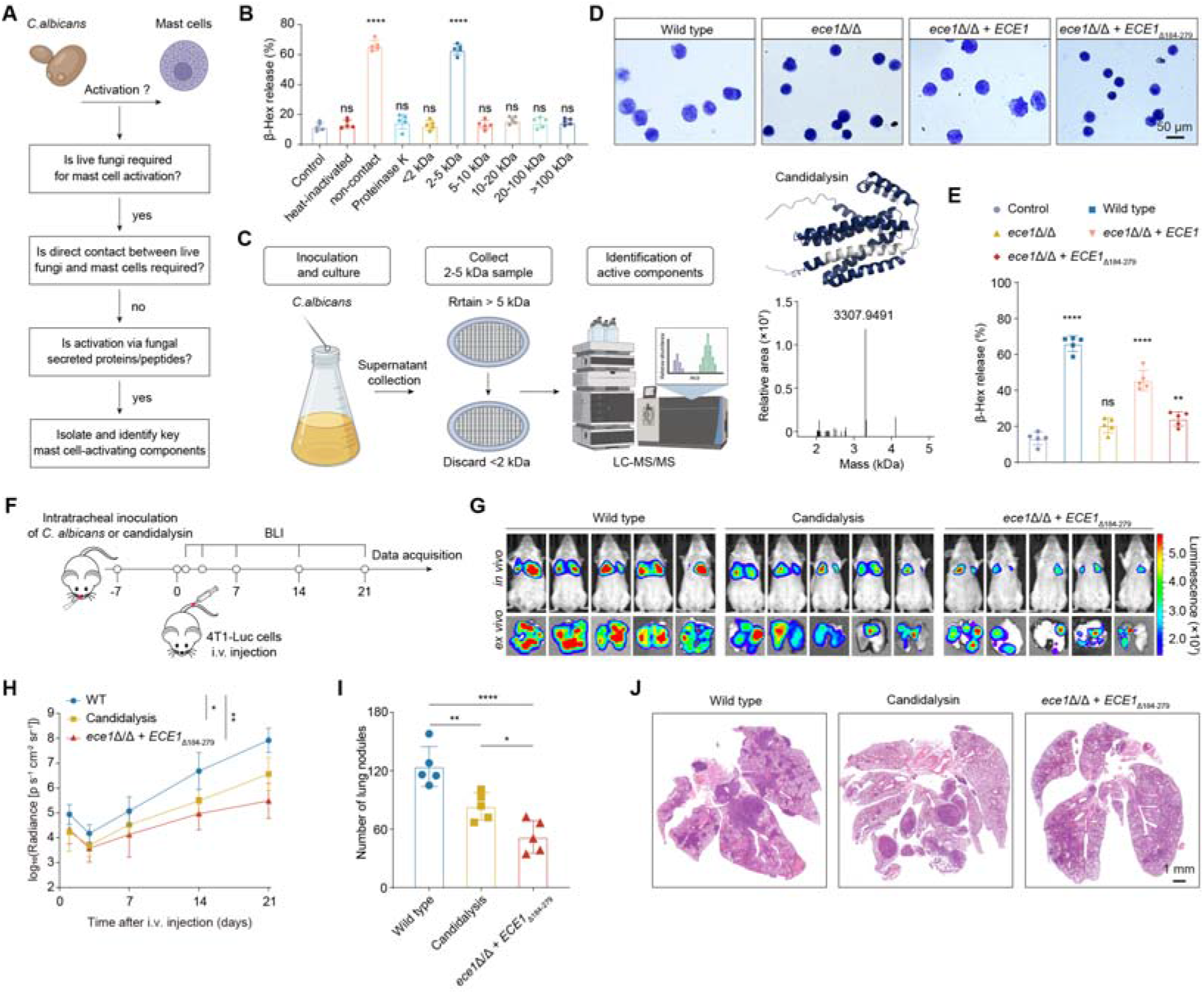
Candidalysin is the key *C. albicans* effector that activates mast cells and drives breast cancer lung metastasis. (A) Schematic of the experimental workflow for identifying the key effector component of *C. albicans* responsible for mast cell activation. (B) Quantification of mast cell degranulation by measuring β-Hex release under the indicated treatment conditions. (C) Identification of candidalysin from active *C. albicans* culture fractions. Schematic of the experimental workflow illustrating supernatant collection, ultrafiltration fractionation, and LC-MS/MS-based identification (left). Predicted three-dimensional structure of candidalysin (top right) and mass spectrometry profile showing a prominent peak at 3307.9491 Da (bottom right). (D–E) Mast cell activation following co-culture with the indicated *C. albicans* strains, assessed by representative toluidine blue staining (D) and quantification of β-Hex release (E). (F) Schematic of the experimental workflow for evaluating the effect of different *C. albicans* strains or candidalysin on breast cancer lung metastasis. (G–H) Representative BLI images of pulmonary metastases (G) and longitudinal bioluminescence tracking of mice (H). (I–J) Quantification of pulmonary metastatic nodules (I) and representative H&E-stained images (J). Data are presented as mean ± SEM. ns, not significant; *p < 0.05; **p < 0.01; ****p < 0.0001 compared using one-way ANOVA (B, E, and H-I).

To test whether candidalysin is required for mast cell activation, mast cells were stimulated with isogenic *C. albicans* strains carrying defined alterations in *ECE1.*^56,57^ Wild-type *C. albicans* and the *ECE1*-complemented strain (*ece1*Δ/Δ *+ ECE1*) induced marked mast cell enlargement and extensive extrusion of metachromatic granules. By contrast, deletion of *ECE1* (*ece1*Δ/Δ) or the candidalysin-encoding domain (*ece1*Δ/Δ *+ ECE*1Δ_184-279_) abolished these morphological changes (Figure 4D). β-hex release assays confirmed that both *ece1*Δ/Δ and *ece1*Δ/Δ *+ ECE*1Δ_184-279_ failed to induce significant mast cell degranulation, whereas complementation with *ECE1* restored this activity (Figure 4E). Thus, *ECE1*-encoded candidalysin is the core fungal effector that specifically triggers mast cell activation.

### Candidalysin-mediated mast cell activation promotes metastatic colonization

We next sought to determine whether candidalysin functionally drives breast cancer lung metastasis *in vivo*. Mice colonized with wild-type *C. albicans*, the candidalysin-deficient mutant, or exogenous candidalysin were challenged intravenously with 4T1-Luc cells (Figure 4F). ^58^ Bioluminescence imaging revealed that mice inoculated with the candidalysin-deficient mutant had significantly lower pulmonary metastatic signals than those colonized with wild-type or receiving exogenous candidalysin (Figures 4G-4H). Endpoint quantification of pulmonary metastatic nodules and H&E staining further confirmed that wild-type colonization and exogenous candidalysin administration increased pulmonary metastatic burden, whereas candidalysin loss markedly abolished this effect (Figures 4I-4J). These data establish candidalysin as a key pro-metastatic fungal effector.

### Candidalysin drives pulmonary mast cell activation and pro-metastatic immune remodeling

Given the potent pro-metastatic effect of candidalysin, we next sought to dissect the immune landscape orchestrated by this axis. We compared the immune microenvironment of lungs colonized with wild-type *C. albicans* versus the candidalysin-deficient mutant *ece1*Δ/Δ *+ ECE*1Δ_184-279_. Flow cytometry and histology showed that wild-type *C. albicans* colonization strongly induced pulmonary mast cell enrichment and activation, whereas the candidalysin-deficient mutant failed to elicit these responses (Figures 5A-5D). Single-cell RNA sequencing (scRNA-seq) further revealed that candidalysin deficiency prevented the expansion and phenotypic remodeling of pulmonary mast cell populations observed following wild-type *C. albicans* colonization (Figures 5E-5F). Avidin-based fluorescence staining showed extensive mast cell degranulation in wild-type-colonized lungs, whereas the proportion of degranulating mast cells was markedly reduced in the mutant group (Figures 5G-5H). These data indicate that candidalysin drives pathological activation of pulmonary mast cells.

**Figure 5.**
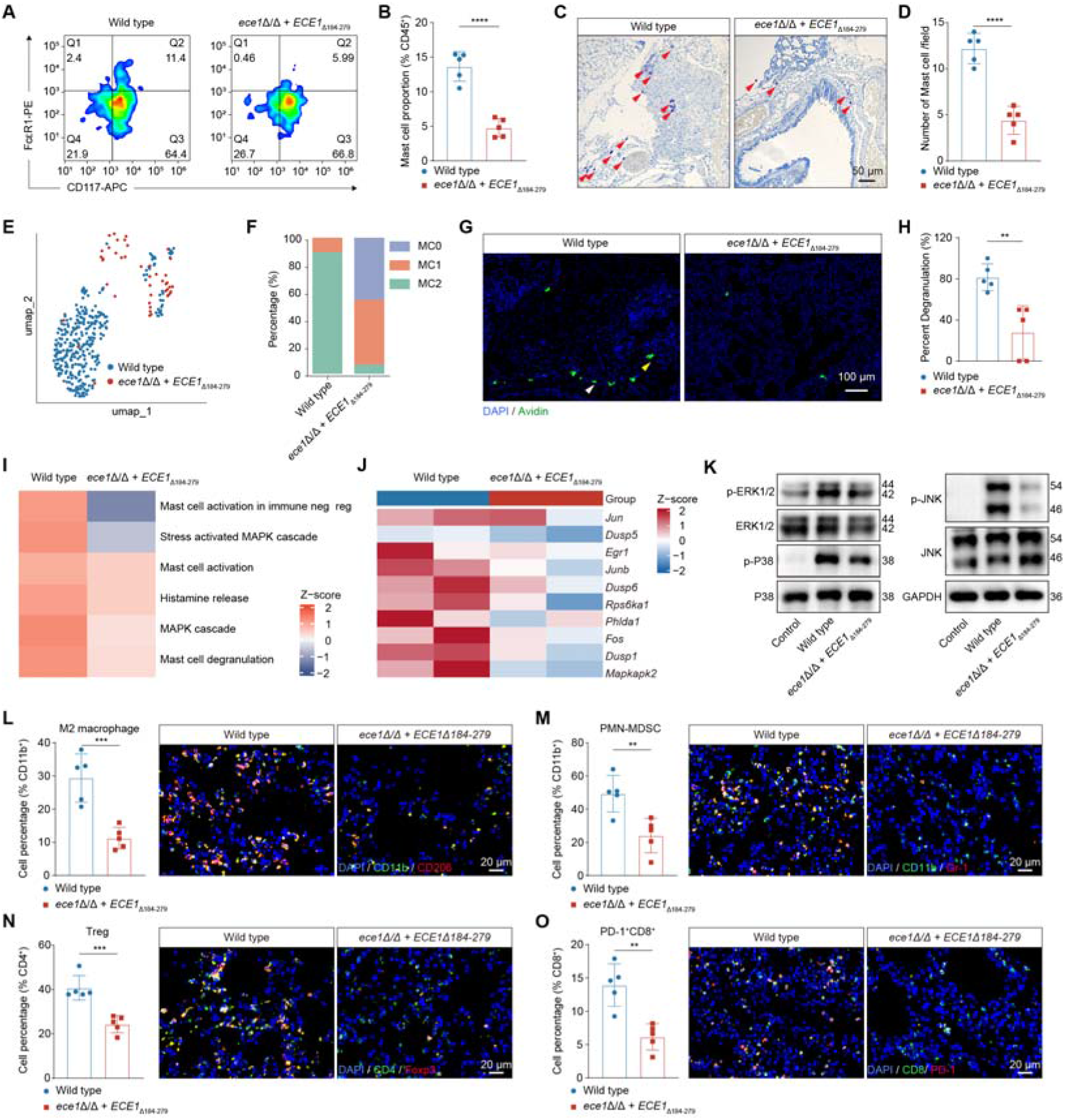
Candidalysin orchestrates pulmonary mast cell activation and immunosuppressive niche formation. (A–B) Mast cell proportions following exposure to wild-type and mutant *C. albicans*. (A) Representative flow cytometry plots. (B) Quantification of mast cell percentages among CD45^+^ cells. (C–D) Representative toluidine blue staining (C) and quantification of mast cells per field (D) in lung tissues. (E–F) UMAP visualization (E) and compositional profiling (F) of pulmonary mast cell populations. (G–H) Representative immunofluorescence images (G) and quantification (H) of degranulated mast cells in lung tissues. DAPI (blue) was used to counterstain nuclei, and Avidin (green) labeled mast cells. Yellow arrows indicate degranulated mast cells; white arrows indicate non-degranulated mast cells. Scale bar, 20 μm. (I–K) Activation of mast cell degranulation and MAPK signaling. (I) Z-score heatmap of pathway activities associated with mast cell degranulation, histamine release, and the MAPK signaling cascade. (J) Heatmap of key effector genes in the MAPK pathway. (K) Western blot analysis of phosphorylated and total ERK1/2, JNK, and P38 protein levels, with GAPDH serving as the loading control. (L–O) Flow cytometric quantification and representative immunofluorescence images of infiltrating immune cell subsets in mouse lungs, showing (L) M2 macrophages, (M) PMN-MDSCs, (N) Tregs, and (O) PD-1^+^ CD8^+^ T cells. Data are presented as mean ± SEM. **p < 0.01; ***p < 0.001; ****p < 0.0001 compared using unpaired Student’s *t* test (two-tailed) (B, D, H, and L-O).

### MAPK signaling axis mediates candidalysin-induced mast cell reprogramming

To dissect the intracellular signaling hub, we performed single-cell transcriptomes and found that mast cells exposed to candidalysin deficient mutant exhibited widespread suppression of inflammatory and immune activation-associated transcriptional programs (Figures S5B-S5D). Wild-type *C. albicans* colonization induced transcriptional programs associated with mast cell degranulation, histamine release, and MAPK signaling, whereas candidalysin deficiency markedly diminished these responses alongside the phosphorylation of core MAPKs, including ERK1/2, JNK, and p38 (Figures 5I-5K).^59,60^ Together, these data demonstrate that candidalysin drives mast cell functional reprogramming through the MAPK pathway.

### Candidalysin reshapes intercellular communication networks in the lung

Because mast cells are key regulators of microenvironment remodeling, we asked whether candidalysin-induced activation shapes the lung into a pro-metastatic niche through intercellular communication. CellChat analysis revealed that the wild-type strain drove dense signaling networks between mast cells and myeloid populations such as macrophages and monocytes, and delivered strong signals to lymphoid cells. Conversely, the candidalysin-deficient mutant substantially weakened these communication networks (Figures S5E-S5F). Quantification of immune composition confirmed that candidalysin deficiency preserved physiological homeostasis of major pulmonary immune populations (Figure S5G).

Consistently, in depth phenotypic profiling showed that wild-type colonization was associated with increased accumulation of immunosuppressive populations, including macrophages (Figure 5L), PMN-MDSCs (Figure 5M), Tregs (Figure 5N), and PD-1 CD8 T cells (Figure 5O). The mutant completely blocked these infiltrations. Consistent with these findings, functional and cell-state signature scoring revealed a marked attenuation of the overall immunosuppressive state in the mutant group (Figure S5H), establishing candidalysin as a key driver of the immunosuppressive pulmonary microenvironment.

### Clinical *C. albicans* isolates exhibit marked heterogeneity in candidalysin secretion

Having established candidalysin as a key driver of *C. albicans*-promoted breast cancer lung metastasis in experimental models, we next assessed its clinical relevance by profiling candidalysin secretion across 279 clinical *C. albicans* isolates (Figure 6A). This analysis uncovered substantial heterogeneity (Figure 6B), with Candidalysin secretion demonstrating a high coefficient of variation (*C.V.* = 1.99) across clinical isolates, indicating that production represents a strain-specific virulence trait rather than a conserved species-level feature. This functional diversity delineated a phenotypic landscape of clinical isolates.

**Figure 6.**
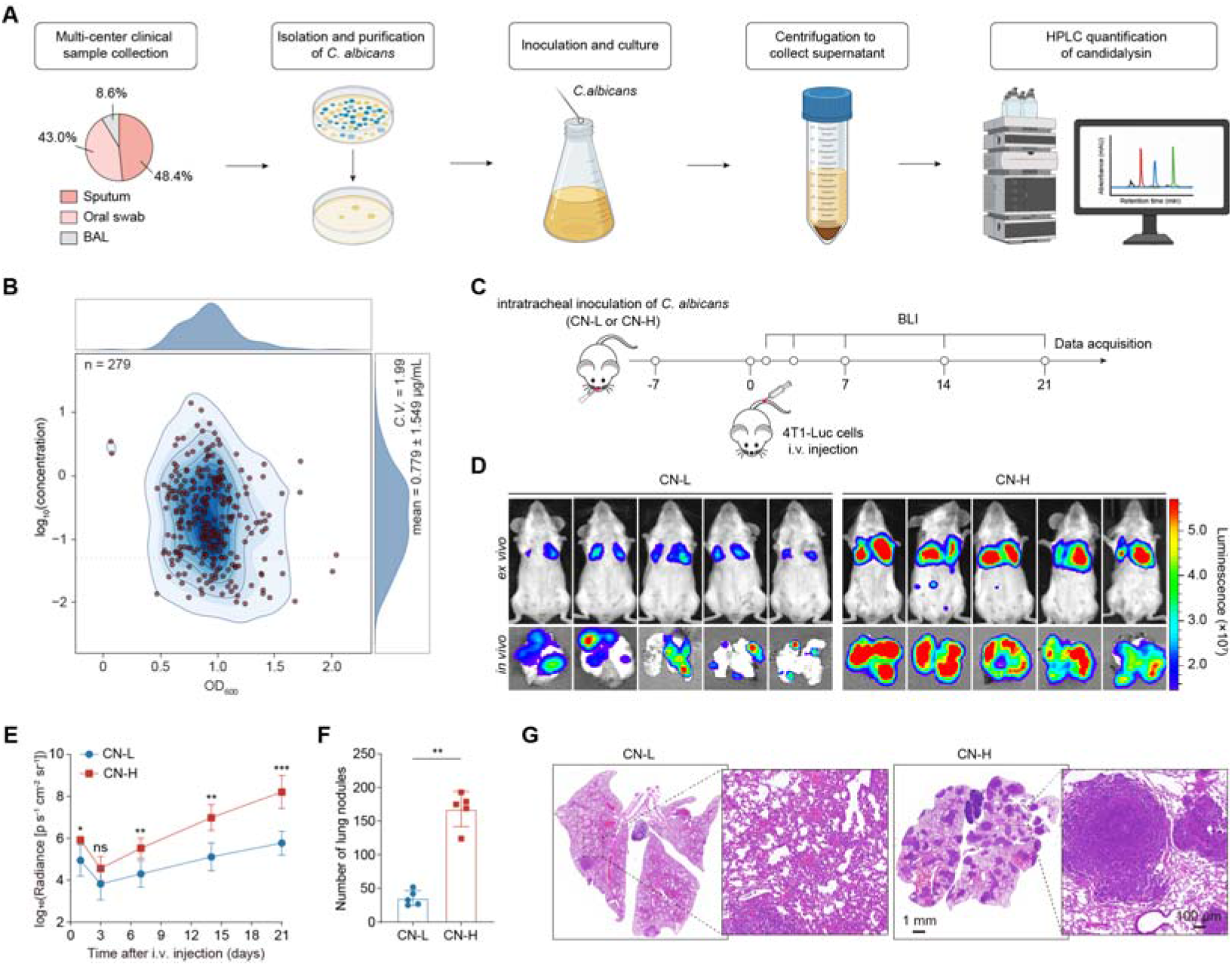
Clinical *C. albicans* isolates with high candidalysin secretion exhibit enhanced pro-metastatic activity. (A) Schematic of the workflow designed for quantitative profiling of candidalysin secretion across clinical *C. albicans* isolates. (B) Candidalysin secretion profiling. Distribution of secretion levels across 279 clinical *C. albicans* isolates, showing log-transformed candidalysin concentration plotted against fungal growth density (OD_600_). (C) Schematic of the experimental workflow evaluating the effects of low-secreting (CN-L) versus high-secreting (CN-H) clinical isolates on breast cancer lung metastasis. (D–E) Representative BLI images of pulmonary metastases (D) and longitudinal bioluminescence tracking of mice (E). (F–G) Quantification of pulmonary metastatic nodules (F) and representative H&E-stained images (G). Data are presented as mean ± SEM. ns, not significant; *p < 0.05; **p < 0.01; ***p < 0.001 compared using unpaired Student’s t test (two-tailed) (E, and G).

### Candidalysin secretion levels determine the metastasis-promoting capacity of clinical isolates in mice

Because candidalysin levels varied by nearly four orders of magnitude, we selected high-secreting (designated CN-H) and low-secreting (designated CN-L) clinical strains. *In vivo*, CN-H-colonized mice exhibited markedly stronger pulmonary bioluminescent signals than CN-L-colonized mice (Figures 6C-6E). Consistently, endpoint nodule counts and histopathology showed higher metastatic incidence and larger areas in the CN-H group (Figures 6F-6G). These results suggest that monitoring clinical fungal secretion profiles may serve as a potential biomarker for breast cancer prognosis and risk stratification.

### Persistent pulmonary immunosuppression limits antifungal and mast cell-stabilizing therapies

In light of the critical roles of *C. albicans* and mast cells in promoting breast cancer lung metastasis, we evaluated the therapeutic efficacy of the mast cell stabilizer cromolyn sodium and the antifungal drug AmBisome under pulmonary *C. albicans* colonization (Figure S6A). Monotherapy failed to suppress metastasis, and combination therapy exerted only limited inhibition (Figures S6B-S6E). Further analysis of the pulmonary tumor microenvironment indicated that, despite engagement of their respective targets, neither treatment reversed the immunosuppressive microenvironment, thereby limiting overall efficacy (Figure S6F).

### Co-encapsulated liposomes synergistically suppress *C. albicans*-mediated lung metastasis

Rg3, a bioactive ginsenoside with well-established immunomodulatory properties,^61,62^ attenuates mast-cell-mediated inflammation by suppressing histamine and pro-inflammatory cytokine release.^63–65^ To achieve coordinated antifungal activity, mast cell stabilization, and reversal of immunosuppression, a liposomal co-delivery system (AG Lip) encapsulating AmB and Rg3 was developed, inspired by the synergistic co-delivery paradigm of the clinically approved liposomal formulation Vyxeos (Figures S6G-S6I). ^66,67^ AG Lip was designed to provide potent antifungal activity together with mast cell-stabilizing effects (Figures S6J-S6O).

AG Lip was next evaluated against *C. albicans*-driven breast cancer lung metastasis (Figure 7A). Of note, AG Lip-treated mice exhibited markedly reduced primary tumor invasiveness, significant decreases in pulmonary metastatic nodule number and volume (Figures 7B-7E), and prolonged survival (Figure 7F). These results demonstrate that AG Lip effectively suppresses *C. albicans* colonization-mediated lung metastatic progression.

**Figure 7.**
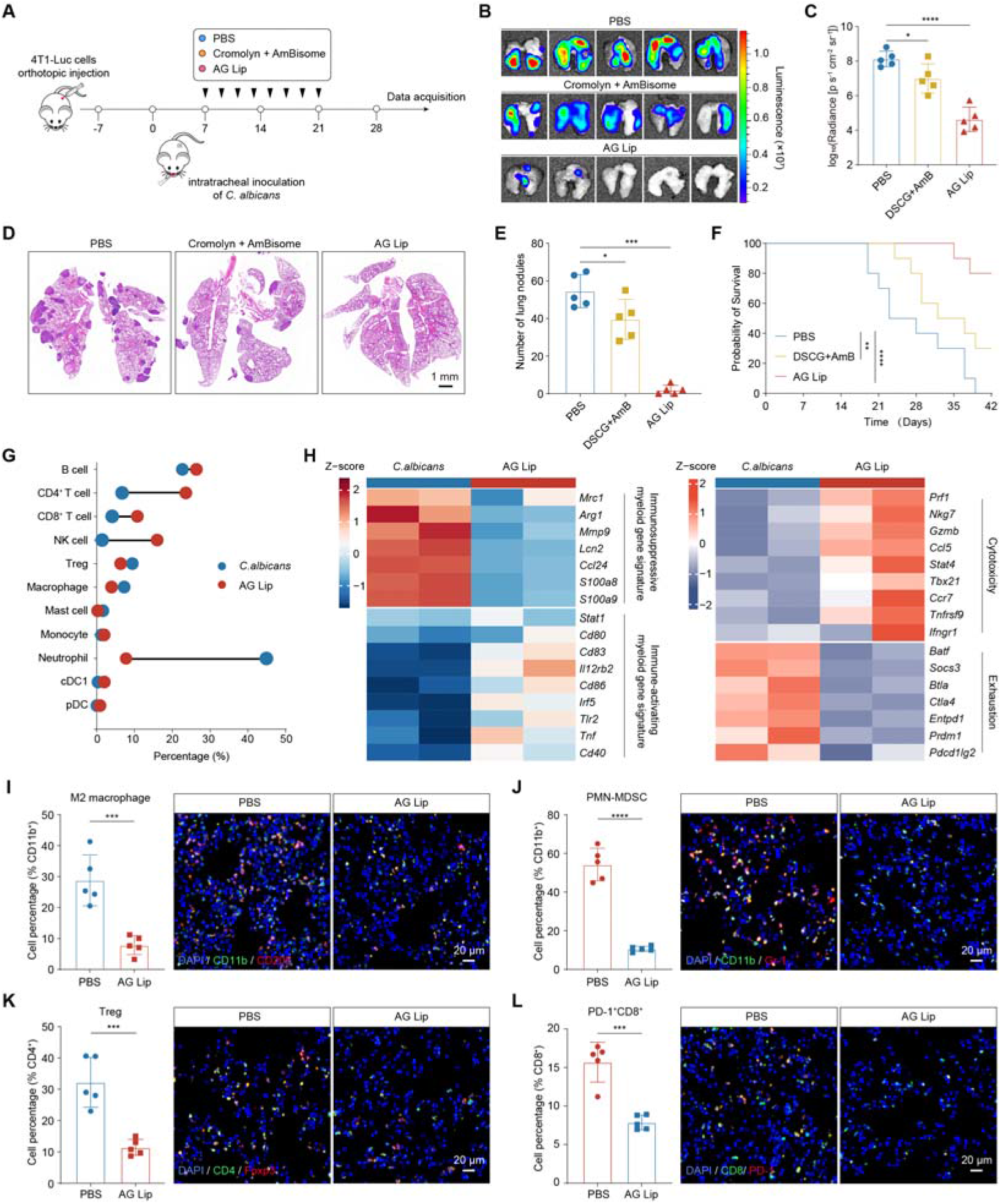
Co-encapsulated liposomes reverse immunosuppression and block fungus-promoted breast cancer metastasis. (A) Schematic of the mouse model and treatment regimen. (B–C) Representative BLI images of pulmonary metastases (B) and longitudinal bioluminescence tracking of mice (C). (D–E) Quantification of pulmonary metastatic nodules (D) and representative H&E-stained images (E). (F) Kaplan-Meier survival curves of mice in the indicated treatment groups. (G) Dumbbell plot showing the proportional shifts of major immune cell subsets in mice. (H) Z-score heatmap of differentially expressed genes (DEGs) in pulmonary myeloid cells and lymphocytes. (I–L) Flow cytometric quantification and representative immunofluorescence images of infiltrating immune cell subsets in mouse lungs, showing (I) M2 macrophages, (J) PMN-MDSCs, (K) Tregs, and (L) PD-1^+^ CD8^+^ T cells. Data are presented as mean ± SEM. *p < 0.05; **p < 0.01; ***p < 0.001; ****p < 0.0001 compared using unpaired Student’s *t* test (two-tailed) (I-L), one-way ANOVA (C and E), and log rank (Mantel-Cox) test (F).

### Co-encapsulated liposomes reshape the immune microenvironment and elicit antitumor immunity

To investigate whether AG Lip remodels the pulmonary immune microenvironment, we performed single-cell RNA sequencing, which revealed that AG Lip treatment significantly reduced pulmonary neutrophils, mast cells, MDSCs, and Tregs, while increasing CD4 T cells, CD8 cytotoxic T cells, and NK cells (Figure 7G; Figures S7A-S7D). Transcriptomic signatures and functional pathway enrichment analyses demonstrated that AG Lip downregulated myeloid-associated immunosuppressive genes, upregulated lymphocyte cytotoxicity-related genes, and promoted pathways related to antigen processing and presentation as well as IFN-γ production (Figure 7H; Figures S7E-S7F). These transcriptional shifts culminated in a marked decrease in lymphocyte exhaustion scores alongside a significant elevation in cytotoxicity scores (Figure S7G), marking a transition of the microenvironment toward an immune-activated state.

Histochemical staining and multiplex immunofluorescence validations further confirmed that AG Lip effectively reduced pulmonary mast cell burden and suppressed their degranulation percentage (Figures S7H-7K), while significantly decreasing the proportions of M2 macrophages, PMN-MDSCs, Tregs, and PD-1 CD8 T cells, thereby dismantling the fungal-induced immunosuppressive network and reconstructing an antitumor immune microenvironment (Figures 7I-7L).

Given this immune reprogramming, AG Lip was further evaluated in combination with the PD-L1-blocking antibody atezolizumab in mice with established lung metastases.^68^ The combination yielded potent synergistic antitumor effects, significantly restraining metastatic progression (Figures S7L-S7Q). These findings indicate that AG Lip dismantles the *C. albicans*-induced immunosuppressive network and activates effector cytotoxic programs, and creates a pulmonary immune context more permissive to checkpoint blockade.

## Discussion

Interactions between the microbiome and the host immune system have become a cornerstone of our understanding of cancer progression.^69–72^ Previous studies have primarily focused on how intratumoral microbiota influences the local tumor microenvironment.^21,30,73^ In contrast, our findings establish that lung-resident *C. albicans* is a key driver of breast cancer lung metastasis, overcoming previous spatial limitations. Our results highlight that the pulmonary mycobiota is not a passive bystander during metastasis. Rather, before tumor cells colonize the lung, it converts this distant organ into a highly permissive pre-metastatic niche through a host-fungus signaling network. This finding expands the classical “seed and soil” hypothesis by incorporating the microbiota into the regulatory network that shapes the pre-metastatic niche.^12,74^ It also advances our understanding of organ-specific metastasis and highlights the importance of monitoring and managing fungal communities as part of cancer therapy.

Having established that pulmonary *C. albicans* colonization remodels the pre-metastatic niche, we next sought to define the effector molecule responsible. We found that candidalysin,^55,56^ a key fungal virulence factor, drives mast cell activation via the MAPK signaling pathway.^56,59^ Activated mast cells release cytokines and chemokines that recruit multiple immunosuppressive populations, including tumor-associated macrophages, myeloid-derived suppressor cells, and Tregs,^44,75^ while inducing functional exhaustion of cytotoxic killer cells. This establishes a highly immunosuppressive pulmonary pre-metastatic microenvironment, disrupting immune homeostasis in the distant organ and providing a molecular explanation for how the fungal community promotes tumor metastasis. However, our data captured only the pre-metastatic stage; the dynamic evolution of this immune landscape during subsequent metastatic stages remains to be defined.

Notably, clinical isolates displayed significant heterogeneity in candidalysin secretion. Higher secretion levels were strongly associated with increased risk of lung metastasis from breast cancer, highlighting its potential as a predictive biomarker for metastatic propensity. These findings suggest that dynamic monitoring of fungal burden and phenotypic characteristics may provide a critical strategy for proactive metastasis prevention in precision oncology. Nevertheless, because of the limited availability of longitudinal clinical cohorts, the relationship between candidalysin secretion capacity and metastatic risk has not yet been validated prospectively in patients; future cohort studies will be needed to establish the generalizability of this mechanism. In addition, the absence of clinically established thresholds for fungal colonization and toxin concentrations that drive tumor metastasis, together with a current lack of highly sensitive detection approaches, limits the translation of these findings into predictive biomarkers or clinical interventions.

Building on this pro-metastatic axis, which links fungal pathogenicity, mast cell hyperactivation, and immune suppression, we developed a liposomal co-delivery system encapsulating AmB and ginsenoside Rg3. This system was designed to disrupt the pro-metastatic signaling cascade through a multidimensional intervention combining pathogen eradication, mast cell stabilization, and reversal of immunosuppression. The delivery system effectively reversed pre-metastatic niche formation and blocked *C. albicans-*driven breast cancer lung metastasis.

Beyond breast cancer, our findings raise the possibility that the pre-metastatic niche shaped by pulmonary *C. albicans* is not cancer-type-specific. Other malignancies that metastasize to the lung may exploit a similar fungus-driven immunosuppressive microenvironment, and commensal microbes residing in other distal organs may likewise prime organ-specific niches for metastatic colonization. This work thus provides a conceptual foundation for understanding how microbial communities across different organs contribute to pre-metastatic niche formation, extending the relevance of our findings beyond a single cancer type.

In summary, lung-resident *C. albicans* actively establishes an immunosuppressive niche that promotes breast cancer cell colonization of the lung through candidalysin-mediated mast cell reprogramming. We further linked heterogeneity in fungal virulence factor secretion to metastatic risk and developed a liposomal co-delivery platform incorporating AmB and ginsenoside Rg3. These findings redefine the role of microbial communities in metastatic niche establishment and provide a new conceptual framework and therapeutic approach for understanding how pathogens drive malignant progression across organs.

## Resource availability

### Lead contact

Requests for further information and resources should be directed to and will be fulfilled by the lead contact, Chong Li.

### Materials availability

Strains and plasmids generated or used in this study are available from the lead contact upon reasonable request.

### Data and code availability

All raw and processed sequencing data, including single-cell RNA sequencing of lung tissue CD45^+^ cells and bulk ITS sequencing, are deposited in public repositories (see key resources table for accession numbers). This paper does not generate any original code. Any requests for additional information or analysis scripts can be made to the lead contact.

## Acknowledgments

This work was supported by the National Natural Science Foundation of China (82073789 [C.L.], 82402651 [L.C.], 32201086 [F.G.], and 82607772 [H.C.]), Sichuan

Science and Technology Program (2026YFHZ0130 [C.L.]), the Research funds from Southern Medical University (2026LH04 [C.L.]) and Chongqing Natural Science Foundation (CSTB2025NSCQ-GPX1349 [L.C.]). We acknowledge Dr. Huan Zhao and Xiaogang Wang (Revvity) for their input on experimental design. We thank Dr. Ningning Liu for providing the *C. albicans* strains associated with Ece1, as well as Dr. Julian Naglik and Dr. Bernhard Hube (original developers of these strains) for making them available for this study.

## Author contributions

C.L., L.W., P.T., L.C., and Z.M. conceived and designed the study. J.W., H.C., Y.T., Y.W., M.R., Y.L., C.C., Y.Z., L.Z., and M.G. performed the investigation and data collection. L.C., C.C., Y.Z., and L.W. provided fungal cultivation, facilities, and technical guidance. Z.M., Y.F., F.G., H.C., X.Y., and L.L. conducted all the experiments and performed the statistical analysis of the data. Z.M., C.L., F.G., and Y.F. wrote the original draft. C.L., L.W., P.T., L.C., Z.M., F.G., C.C., and L.L. reviewed and edited the manuscript. All authors discussed the experiments and results and approved the manuscript.

## Methods

### Cell lines

Mouse breast cancer cell lines (4T1 and E0771) and HEK 293T cells were obtained from the National Collection of Authenticated Cell Cultures (Chinese Academy of Sciences, Shanghai, China). Cell lines were cultured in DMEM medium (Gibco, C11995500BT) supplemented with 10% fetal bovine serum (FBS; Gibco, 10099-141) and 1% penicillin-streptomycin (Gibco, 15140122) and maintained at 37°C in a humidified incubator containing 5% CO . All cell lines were routinely tested for mycoplasma contamination before and after cryopreservation or experimental use using the MycoBlue Mycoplasma Detection Kit (Vazyme, D101-02) according to the manufacturer’s instructions, and were confirmed to be mycoplasma-free.

To generate reporter cell lines for *in vivo* monitoring of tumor burden, firefly luciferase and ZsGreen fluorescent protein expression cassettes were integrated into the genomes of 4T1 and E0771 cells through lentiviral transduction. Briefly, HEK293T cells were seeded in culture dishes and cultured for 18-24 h until reaching approximately 70% confluence. Cells were then transfected with either the plasmid CMV enhancer-MCS-hEF1/HTLVp-firefly_Luciferase-T2A-Neomycin (Genechem, GV870) or CMV enhancer-MCS-polyA-EF1A-zsGreen-sv40-puromycin (Genechem, GV658). At 48 and 72 h after transfection, culture supernatants containing lentiviral particles were collected and concentrated by ultracentrifugation at 20,000 rpm for 2 h. The concentrated lentiviral preparations were subsequently used to transduce 4T1 and E0771 cells. Following transduction, cells were continuously selected with the corresponding antibiotics to establish stable luciferase(luc)- or ZsGreen(ZsG)- expressing cell lines, which were subsequently used for cell expansion and in vivo imaging studies.

### Fungal strains and culture conditions

The fungal strains used in mouse experiments are listed in Table S3. All *C. albicans* strains associated with Ece1 used in this study were kindly provided by Dr. Ningning Liu, with explicit permission from the original developers of these strains, Dr. Julian Naglik and Dr. Bernhard Hube. All strains were stored at -80°C. Before each experiment, strains were recovered in YEPD medium (1% yeast extract, 2% peptone, and 2% glucose) and cultured at 30°C for 10-12 h. Fungal cells were then harvested, washed three times with PBS.

### Animals

All animal experiments were performed in accordance with institutional guidelines and were approved by the IACUC of Southern Medical University (Approval No. 202412013) and Southwest University (Approval No. 20240109-06). BALB/c and C57BL/6 mice were purchased from SJA Laboratory Animal Co., Ltd. (Changsha, China). *Cpa3*-2A-CreERT2 mice (NM-KI-200005) were obtained from Shanghai Model Organisms Center, Inc., and The *ROSA26*-eGFP-DTA mice (RRID: IMSR_JAX:006331) were obtained from The Jackson Laboratory.^76^ *Cpa3*-2A-CreERT2 mice were crossed with *ROSA26*-eGFP-DTA mice to generate mast cell-conditional ablation mice (*Cpa3*-CreERT2 / ; *ROSA26*-DTA / mice). Genotypes were determined by PCR using a mouse tail genotyping kit (Beyotime, D7283S) (primer sequences listed in the Key Resources Table).

### Human patient samples

Frozen specimens (n = 12, 29, and 18 for biopsy samples, surgical resection samples, and BFA, respectively), fresh specimens (n = 5, 6, and 12 for biopsy samples, surgical resection samples, and BFA, respectively), formalin-fixed paraffin-embedded (FFPE) specimens (n = 16 and 59 for biopsy samples and surgical resection samples, respectively), and follow-up information (n = 66 for overall survival analysis, respectively) were obtained from the First Affiliated Hospital of Army Medical University (IRB-approved protocols: KY2023144), the Chongqing General Hospital (IRB-approved protocols: S2025-052-01), the First Affiliated Hospital of Xi’an Jiaotong University (IRB-approved protocols: XJTU1AF2026LSYY-0189), the Chongqing University Cancer Hospital (IRB-approved protocols: CZLL2026-162-001), and the Second Affiliated Hospital of Chengdu Medical College (IRB-approved protocols: KJ2022025). Clinical *C. albicans* isolates (n = 135, 120, and 24 for sputum, oral swabs, and BFA, respectively) were obtained from the Huashan Hospital, Fudan University (IRB-approved protocol: KY2026-081), the First Affiliated Hospital of Army Medical University, the Chongqing General Hospital, and the Second Affiliated Hospital of Chengdu Medical College. All participants provided written informed consent, and all procedures involving human samples were approved by the institutional ethics committees of the participating hospitals.

### Fungal DNA extraction

Appropriate amounts of sample material were transferred into microcentrifuge tubes containing extraction lysis buffer and mechanically disrupted by grinding. Genomic DNA was subsequently extracted using the MagBeads FastDNA Kit for Soil (MP Biomedicals, 116564384) according to the manufacturer’s instructions. The extracted DNA was evaluated by 0.8% agarose gel electrophoresis to assess DNA integrity and fragment size. DNA concentration and purity were determined using a NanoDrop spectrophotometer (Thermo Scientific, NC2000).

### ITS1 rRNA sequencing and analysis

High-throughput sequencing of the fungal internal transcribed spacer 1 (ITS1) rRNA region was performed on an Illumina NovaSeq platform. Sequence data analyses were mainly performed using QIIME2 and R packages (v3.6.0).^77^ Briefly, raw sequencing data were demultiplexed using the demux plugin, and primer sequences were removed using the cutadapt plugin.^78^ Subsequently, the DADA2 plugin was applied for quality filtering, denoising, paired-end read merging, and chimera removal.^79^ The resulting high-quality sequences were clustered into amplicon sequence variants (ASVs) at 100% sequence identity, enabling single-nucleotide resolution profiling, and an ASV abundance table was generated. Representative ASV sequences were taxonomically classified by comparison against the UNITE database. ASV abundance matrices containing rare ASVs were removed before downstream analyses.

### Fluorescence in situ hybridization (FISH) analysis

Paraffin-embedded tissue sections obtained from FFPE blocks were washed in 0.1 M Tris-HCl buffer (pH 7.4) for 15 min and treated with lysozyme (10 mg/mL) for 30 min to enzymatically digest the fungal cell wall. Subsequently, sections were hybridized with a *C. albicans*-specific 23S rRNA probe labeled with Cy3 at the 5′ end (sequence: ACAGCAGAAGCCGTGCC),^80^ which targets fungal 23S ribosomal RNA (rRNA) localized within the fungal cytoplasm. Fungal colonization in breast cancer lung metastatic tissues was visualized by FISH using the Pannoramic MIDI digital slide scanning system.

### Tumor models

For the orthotopic 4T1 tumor model, 4T1 cells (5 × 10^5^ cells per mouse) were injected into the fourth mammary fat pad on the left side of BALB/c mice. Seven days after tumor implantation, when palpable tumors were established, mice were randomly assigned to different experimental groups and inoculated with fungal strains via intratracheal administration. Tumor-bearing mice were monitored regularly, and tumor volumes were measured every three days using calipers according to the following formula: Volume = (tumor length) × (tumor width)^2^/2. Mice were monitored for clinical signs, including reduced food intake, body weight loss, or dehydration. For the lung metastasis model, mice were first inoculated with fungal strains via intratracheal administration. Seven days later, 4T1-Luc cells (5 × 10 cells per mouse) were injected intravenously (*i.v.*) through the tail vein in 200 μL HBSS (Biosharp, BL559A).

### Intratracheal fungal inoculation and optimization of *C. albicans* dosage

Mice were anesthetized by intraperitoneal injection of pentobarbital sodium (60 mg/kg body weight). After anesthesia was confirmed by regular thoracoabdominal respiration, muscle relaxation, and absence of withdrawal reflexes, mice were placed on an intubation platform. The oral cavity was gently opened to expose the oropharynx. Mice under anesthesia with spontaneous respiration were intratracheally administered 50 μL of PBS (Biosharp, BL302A) containing graded doses of *C. albicans* cells (1 × 10^4^, 1 × 10 , 1 × 10^6^, 1 × 10^7^, or 1 × 10^8^ cells).^81,82^ Following intratracheal inoculation, mice were monitored for clinical signs, including reduced food intake, body weight loss, or dehydration, and survival rates were recorded. Animals were euthanized when body weight loss reached 20%, and were considered deceased for survival analysis.

### Bioluminescence imaging (BLI)

For metastasis studies, whole-body *in vivo* imaging and *ex vivo* organ bioluminescence imaging were performed using an IVIS imaging system to assess metastatic burden. Mice were anesthetized and intraperitoneally injected with D-luciferin (Gold Biotechnology, LUCK-1G, 150 mg/kg).^83^ After 5 min, bioluminescence signals were acquired. For *ex vivo* imaging, tissues were collected after PBS perfusion and incubated in 1× PBS containing D-luciferin (15 mg/mL) at room temperature for at least 15 min to allow complete substrate penetration. Bioluminescence signals were acquired using an IVIS Lumina Series III imaging system with an exposure time of 1 min, and image acquisition and quantitative analysis were performed using Living Image software (Revvity).

### Histopathological Analysis

Lung tissues were harvested and fixed in 4% paraformaldehyde (PFA) at 4°C overnight, followed by three washes in PBS. Surface metastatic nodules on the lungs were quantified manually under a stereomicroscope (RWD, 77001S). Subsequently, individual lung lobes were microdissected from the trachea, routinely processed, and co-embedded in paraffin blocks for complete cross-sectional sectioning at a thickness of 3.5 μm. Histological damage and inflammatory cell infiltration were evaluated by Hematoxylin and Eosin (H&E) staining. Metastatic burden was quantified using ImageJ software (NIH) and expressed as the percentage of metastatic lesion area relative to the total evaluated lung tissue area (%). *C. albicans* pulmonary colonization was assessed via Periodic Acid-Schiff (PAS) staining. Mast cell infiltration was examined using toluidine blue staining.

### Determination of the pre-metastatic stage in the breast cancer model

To define the pre-metastatic and metastatic phases in the 4T1 breast cancer model, we performed flow cytometric and fluorescence imaging analyses. Briefly, mice were orthotopically implanted with 4T1-ZsG cells (5 × 10 cells per mouse), and the proportion of ZsGreen-positive cells in lung tissues was quantified at days 7, 14, and 21 post-tumor implantation. Primary tumor tissues and lung samples from naive mice (day 0) served as positive and negative controls, respectively.^84^ The pre-metastatic stage was defined as the phase preceding the emergence of overt metastatic lesions in distant lung tissues, characterized by undetectable or minimal tumor cells in the lungs. Based on these analyses, the pre-metastatic window in the 4T1 model was established as occurring on or before day 14.

### Mast cell differentiation and culture

Mouse mast cells were generated from bone marrow cells isolated from C57BL/6 mice. Bone marrow cells were cultured in DMEM supplemented with 10% FBS, 1% penicillin-streptomycin, 2 mM β-mercaptoethanol (MCE, HY-Y0326), with stimulation by stem cell factor (SCF, BioWorld, BK0345, 20 ng mL ¹) and interleukin-3 (IL-3, BioWorld, BK0103, 10 ng mL ¹) for 2-4 weeks to induce mast cell differentiation.^85,86^ After 48 h of culture, non-adherent cells were transferred to new culture flasks, and the medium was subsequently replaced weekly to remove adherent cells. For *in vitro* experiments, compounds were used according to the manufacturer’s recommended instructions. For *in vivo* experiments, mast cells (CD117 FcεRI ; purity ≥95%) were isolated by flow cytometric sorting from orthotopic tumors of mice with the indicated genetic backgrounds and used within 4 h to maintain cell viability.

### Tamoxifen-induced Cre recombination

To induce Cre-mediated recombination in *Cpa3*-CreERT2 / ; *ROSA26*-DTA / mice *in vivo*, tamoxifen (Ron Reagent, R019059) was dissolved in corn oil at a concentration of 20 mg/mL, incubated at 37°C for 2 h, and further sonicated to facilitate complete dissolution. Female *Cpa3*-CreERT2 / ; *ROSA26*-DTA / mice were administered tamoxifen intraperitoneally (120 mg/kg) once every 24 h for five consecutive days starting at 6-8 weeks of age.^87,88^ Seven days after the final tamoxifen injection, mice were intratracheally inoculated with fungal strains, followed by intravenous injection of E0771-Luc cells via the tail vein 7 days after fungal inoculation. To maintain sustained recombinase activity, mice received additional intraperitoneal injections of tamoxifen (100 μL) every 4 days.

### Cytokine measurement

Cytokine levels in cell culture supernatants or serum samples were quantified using ELISA kits according to the manufacturer’s instructions. Mouse CXCL2 (Mlbio, ML058180), mouse IL-1β (Mlbio, MIC50300-1), mouse CCL2 (Mlbio, ML037840), mouse CCL3 (Mlbio, ML063493), mouse IL-6 (Mlbio, ML063159) and mouse TNF-α (Mlbio, ML002095) were measured using corresponding ELISA kits. Briefly, 100 μL of cell culture supernatant was added to each pre-coated well and incubated for 2 h at room temperature. After washing three times with washing buffer, biotin-conjugated detection antibodies diluted in assay diluent were added and incubated for 2 h at room temperature. After three additional washes, HRP-conjugated streptavidin was added and incubated for 45 min at room temperature. Finally, TMB substrate solution was applied for color development, and absorbance was measured at 450 nm.

### β-Hex release assay

To evaluate mast cell degranulation and granule content release, the release rate of β-Hex was quantitatively measured. Mast cells were counted, resuspended in Tyrode’s buffer (procell, PB180338) containing 2% BSA, and adjusted to a density of 4 × 10 cells/mL. 500-μL aliquot of cell suspension was added to the lower chamber of a 24-well Transwell plate (Corning, 3413) and equilibrated at 37°C with 5% CO for 10 min. Subsequently, 100 μL of *C. albicans* suspension (1 × 10 cells per well) was added to the upper chamber to stimulate mast cell degranulation, while 50 μL of Tyrode’s buffer alone was added to the control group. After incubation at 37°C with 5% CO for 2 h, the contents of the lower chamber were collected into 1.5 mL microcentrifuge tubes and centrifuged at 300 × g for 5 min at 4°C to terminate the degranulation reaction. Cell-free supernatants were collected and kept on ice. The remaining cell pellets were resuspended in 500 μL Triton X-100 lysis buffer (0.5% Triton X-100 prepared in PBS) and lysed on ice for 15 min. Subsequently, 50 μL of cell lysates or degranulation supernatants were transferred into 96-well flat-bottom plates and mixed with 50 μL of reaction solution containing 2 mM 4-nitrophenyl-N-acetyl-β-D-glucosaminide (pNAG, MCE, HY-W011183) dissolved in 0.1 M citrate buffer (pH 4.5). Plates were incubated at 37°C for 1 h. The enzymatic reaction was terminated by adding 100 μL of 0.1 M carbonate buffer (100 mM Na CO and 100 mM NaHCO , pH 9.0) to each well. Absorbance was measured at 405 nm using a microplate reader (Synergy H1, BioTek) to quantify β-Hex activity. The percentage of β-Hex release was calculated as the amount of β-Hex in the supernatant divided by the total β-Hex content (β-Hex in the supernatant plus β-Hex in the cell lysate).

### Cell migration assay

Cell migration was evaluated using a Transwell migration assay (Corning, 3422). Briefly, mast cells were pre-cultured and serum-starved in serum-free DMEM (Gibco, C11995500BT) for 24 h. Subsequently, 2 × 10 cells were seeded into the upper chamber containing serum-free DMEM, while the lower chamber was supplemented with DMEM containing 10% FBS and *C. albicans* to evaluate its effect on mast cell migration. After 24 h of incubation, non-migrated cells remaining on the upper surface of the membrane were gently removed using cotton swabs. Migrated cells on the lower surface of the membrane were fixed with 4% PFA and stained with crystal violet (Beyotime, C0121). The percentage of migrated area in each replicate was quantified using ImageJ software. Each experiment was independently repeated three times.

### Flow cytometric analysis of lung immune cells

For pulmonary immune profiling, 6-8-week-old female mice received an orthotopic injection of 5 × 10 4T1 cells into the left fourth mammary fat pad. Seven days after tumor implantation, mice were intratracheally inoculated with 5 × 10 fungal cells. Fourteen days after fungal inoculation, mice were anesthetized for isolation of pulmonary immune cells. The abdominal and thoracic cavities were carefully opened to expose the heart while avoiding injury to major blood vessels and lung tissues. The abdominal aorta was incised, and the lungs were perfused through the right ventricle until complete blanching was observed, indicating removal of circulating blood. Lungs were then harvested, and single-cell suspensions were generated using the Lung Dissociation Kit, mouse (Miltenyi Biotec, 130-095-927) according to the manufacturer’s instructions. Cell suspensions were filtered through 70 μm sterile cell strainers (Biosharp, BS-70-CS), and the digestion tubes were rinsed with an additional 10 mL DMEM/F12 medium. Cells were centrifuged at 500 × g for 5 min, followed by red blood cell lysis with 5 mL ACK lysis buffer (Beyotime, C3702) on ice for 5 min. Cells were washed once with HBSS (Biosharp, BL559A), centrifuged, and resuspended for immediate flow cytometric analysis.

For lymphoid cell profiling, cells were stained with antibodies against mouse CD45-PE/Cy7 (BioLegend, 103114), CD3-FITC (BioLegend, 100204), CD8a-PE (BioLegend, 100708), CD4-APC (BioLegend, 100412), CD4-APC (BioLegend, 100412), CD4-PerCP/Cy5.5 (BioLegend, 100433), CD25-APC (BioLegend, 102012), Foxp3-PE (BioLegend, 126404), CD279-APC (BioLegend, 135209), Fc block (ABclonal, A29013), and Zombie NIR (BioLegend, 423106). For myeloid cell profiling, cells were stained with antibodies against mouse CD45-PE/Cy7, CD11b-FITC (BioLegend, 101206), F4/80-PerCP (BioLegend, 123126), CD206-APC (BioLegend, 141708), CD86-PE (BioLegend, 105008), Ly6C-APC (BioLegend, 128016), Ly6G-PE (BioLegend, 127608), CD117-APC (BioLegend, 135108), FcεRIα-PE (BioLegend, 134307), Fc block, and Zombie NIR. Flow cytometric analysis was performed using a Beckman CytoFLEX 5L flow cytometer, and cell sorting was performed using a Beckman CytoFLEX SRT cell sorter.

### Immunofluorescence and immunohistochemistry staining

For immunofluorescence staining, paraffin-embedded tissue sections were fixed in 4% PFA at 4°C for 24 h, followed by dehydration through graded ethanol solutions and clearing with xylene before paraffin embedding. Tissue sections were cut using a microtome, deparaffinized, and rehydrated. Antigen retrieval was performed by heating sections in boiling citrate buffer (10 mM sodium citrate, 0.05% Tween-20, pH 6.0) using a microwave oven at medium-low power for 15 min. Sections were blocked in blocking buffer (TBS containing 3% BSA, and 0.1% Triton X-100) for 1 h at room temperature in a humidified chamber. Sections were then incubated with primary antibodies, including anti-CD11b (ServiceBio, GB150266), anti-CD206 (ServiceBio, GB153497), anti-Ly6G (ServiceBio, GB11229), anti-CD4 (ServiceBio, GB15064), anti-Foxp3 (ServiceBio, GB150152), anti-CD8 (ServiceBio, GB15068), and anti-PD-1 (ServiceBio, GB153744). After primary antibody incubation, sections were incubated with corresponding secondary antibodies. Nuclei were counterstained with DAPI (Servicebio, G1012), and slides were mounted using an antifade mounting medium (Servicebio, G1401).

For immunohistochemical staining, paraffin-embedded breast tumor sections were deparaffinized and rehydrated, followed by antigen retrieval, endogenous peroxidase blocking, blocking, and incubation with anti-Ki67 antibody (Servicebio, GB151499). Antigen signals were visualized using a DAB chromogenic substrate (Servicebio, G1212). Sections were subsequently counterstained with hematoxylin, differentiated with acid differentiation solution, dehydrated, and mounted with neutral resin. Images were acquired and analyzed using a digital pathology slide scanner (Pannoramic, 250/MIDI).

### Quantitative real-time PCR analysis

Total RNA was extracted using the RNA extraction kit (Vazyme, RC201) according to the manufacturer’s instructions. RNA was reverse transcribed into cDNA using the HiScript® II Q RT SuperMix for qPCR (+gDNA wiper) kit (Vazyme, R222-01). Primers targeting specific genes were designed and synthesized by Sangon Biotech Co., Ltd. (Shanghai, China). The cDNA samples were subjected to pre-amplification for 15-20 cycles using ChamQ Universal SYBR qPCR Master Mix (Vazyme, Q711-03) on a PCR system. Subsequently, gene-specific quantitative PCR analysis was performed on a Real-Time PCR System (Rocgene, China) according to the manufacturer’s instructions. Data acquisition and analysis were performed using Jena analysis software and Microsoft Excel. Primers used are listed in key resources table.

### Western blotting

Protein lysates were extracted from mast cells using RIPA lysis buffer (Beyotime, P0013B) supplemented with phenylmethylsulfonyl fluoride (PMSF, Beyotime, ST506) and a protease inhibitor cocktail (Sigma, 11836170001). Protein samples were separated by 10% SDS-PAGE and transferred onto polyvinylidene difluoride (PVDF) membranes (Millipore, IPVH00010). PVDF membranes were blocked with 5% BSA in TBST (Servicebio, G2150-1L) for 1 h at room temperature and incubated with primary antibodies overnight at 4°C, followed by incubation with corresponding secondary antibodies for 1 h. After washing with TBST, protein signals were detected using chemiluminescent substrate (Thermo Scientific, Cat#34580) and visualized using a chemiluminescence imaging system (Gel Image System, China). The primary antibodies used were anti-ERK1/2 antibody (Abcam, ab184699), Anti-JNK antibody (Abcam, ab179461), anti-p38 antibody (Huabio, ET1702-65), anti-p-ERK1/2 antibody (Abcam, ab201015), Anti-p-JNK antibody (Beyotime, AB3065), anti-p-p38 antibody (Huabio, HA722669), anti-GAPDH antibody (Proteintech, 10494-1-AP). The secondary antibody was HRP-conjugated anti-rabbit IgG antibody (Proteintech, SA00001-2).

### Preparation of single-cell suspensions

Lung tissues were collected and transferred into RPMI-1640 medium (Gibco, C11875500BT) supplemented with 1% penicillin-streptomycin. Tissues were minced into approximately 0.5 mm³ fragments and enzymatically digested using the MACS Lung Dissociation Kit, mouse (Miltenyi Biotec, 130-095-927), according to the manufacturer’s instructions, with agitation at 37°C for 30 min. The digested cell suspensions were sequentially filtered through 70 μm and 40 μm cell strainers (Biosharp, BS-40-CS, BS-70-CS) and centrifuged at 300 × g for 10 min. Cell pellets were resuspended in ACK Lysis Buffer and incubated on ice for 2 min to remove erythrocytes. After washing twice with PBS, cells were resuspended in PBS containing 0.04% bovine serum albumin (BSA) for subsequent analysis.

### scRNA-seq library construction and sequencing

Single-cell RNA sequencing libraries were generated using the DNBelab C Series high-throughput single-cell RNA library preparation kit (MGI, Cat#940-001818-00). Briefly, single-cell suspensions were encapsulated into droplets, followed by emulsion disruption, mRNA capture using barcoded magnetic beads, reverse transcription, cDNA amplification, and purification to generate barcoded single-cell RNA sequencing libraries. The resulting cDNA products were enzymatically fragmented into 300-500 bp fragments and used for indexed sequencing library construction according to the manufacturer’s instructions. Library quality was assessed using the Qubit ssDNA Assay Kit (Thermo Fisher Scientific) and an Agilent 2100 Bioanalyzer. Qualified libraries were subjected to paired-end sequencing on MGI sequencing platforms, including MGISEQ-2000.

### scRNA-seq data processing

Sequencing data were processed using an open-source analysis pipeline (https://github.com/MGI-tech-bioinformatics/DNBelab_C_Series_scRNA-analysis-so ftware). Briefly, sequencing reads from all samples were demultiplexed, barcode information was processed, and 3′ end unique molecular identifier (UMI) counting was performed using default parameters. Processed sequencing reads were aligned to the human reference genome (GRCh38) using STAR (version 2.5.1b). Valid cell barcodes were automatically identified based on the distribution of UMI counts per cell using the barcodeRanks function implemented in the DropletUtils package. Background droplets and barcodes with UMI counts below the predefined threshold were removed. Gene expression levels were subsequently quantified using the PISA pipeline, and gene-by-cell expression matrices were generated for each sequencing library.

### scRNA-seq data analysis

Raw gene expression matrices (generated by the MGI pipeline) were imported into R (v4.5.0) using the ‘ReadMtx’ function and converted into Seurat (version 5) objects, retaining genes detected in ≥3 cells and cells expressing ≥200 genes. For each sample, low-quality cells with fewer than 300 genes (nFeatureRNA), more than 5,000 genes (nFeatureRNA), or a mitochondrial gene percentage (percent.mt) greater than 8% were discarded. After the first quality control, 1,493,240 cells remained for the subsequent analyses. Surviving cells were log-normalized, and the 2,000 most variable genes were selected using the variance-stabilizing transformation (VST) method. Expression matrices were then normalized to a scale factor of 10,000 transcripts per cell and subsequently centered and scaled prior to dimensionality reduction. Principal component analysis (PCA) was performed on the scaled data using 50 principal components (PCs). To mitigate inter-replicate batch effects, Harmony integration was applied to the PCA embeddings, with "sample" used as the grouping variable, and the first 30 Harmony dimensions were retained for downstream analyses. The batch-corrected embeddings were used for graph-based clustering and Uniform Manifold Approximation and Projection (UMAP) visualization.

### Preparation of AG Lip

A defined amount of egg phosphatidylcholine and ginsenoside Rg3 (Phytopurify, BP0040) was dissolved in an appropriate volume of anhydrous ethanol and transferred into a clean, dry round-bottom flask. The organic solvent was removed by rotary evaporation under reduced pressure until a thin lipid film was formed. Amphotericin B (AmB, MCE, HY-B0221) was dissolved in 50 μL DMSO (MCE, HY-Y0320C) and mixed with 950 μL of 5% (w/v) glucose solution. The resulting solution was added to the flask, and the flask was gently shaken to detach the lipid film from the inner wall. The resulting suspension was sonicated at 600 W until translucent, then purified to remove unencapsulated AmB, and finally filtered through a 0.22 μm membrane.^89^ Unmodified liposomes (AC Lip) were prepared using the same procedure, except that cholesterol was used instead of ginsenoside Rg3.

### Characterization of liposomes

The hydrodynamic diameter, zeta potential, and polydispersity index (PDI) of AG Lip and AC Lip were measured by dynamic light scattering (DLS) assay (Malvern, UK). The morphology of AG Lip and AC Lip was examined by transmission electron microscopy (TEM; Tecnai G2 spirit, FEI, USA).

### *In vivo* treatment

All treatments were administered on days 7, 9, 11, 13, 15, 17, 19, and 21 after tumor implantation. AmBisome was administered intravenously via tail vein injection at an amphotericin B (AmB) dose of 1 mg/kg. Sodium cromoglycate was administered orally at 20 mg/kg. AG Lip and AC Lip were administered intravenously via tail vein injection at an AmB-equivalent dose of 1 mg/kg. For combination therapy experiments, stable 4T1-Luc cells expressing human PD-L1 (4T1-Luc-hPD-L1) were generated by transfection with plasmid CMV enhancer–MCS–hPD-L1–Puromycin expression vector.^90^ Mice were intratracheally inoculated with *C. albicans*, followed 7 days later by intravenous injection of 4T1-Luc-hPD-L1 cells to establish a pulmonary *C. albicans* colonization-associated breast cancer lung metastasis model.

Tumor-bearing mice with pulmonary *C. albicans* colonization were treated with anti-PD-L1 antibody atezolizumab (2.5mg/kg; MCE, HY-P9904; i.v.) alone or in combination with AG Lip.^91,92^ Detailed dosing information and treatment schedules for each experiment are provided in the corresponding figure legends and schematic diagrams.

### High-performance liquid chromatography analysis

*C. albicans* culture supernatants were filtered using Amicon™ Ultra centrifugal filters with a 10 kDa molecular weight cutoff (Millipore) and subsequently subjected to high-performance liquid chromatography (HPLC) analysis using a Shimadzu system with LabSolutions LC software. Chromatographic separation was performed using a C18 column (Diamonsil, 250 × 4.6 mm, 5 μm, Cat #99603). The major product peaks were identified by comparison with a synthetic candidalysin standard (MCE, HY-P10408). The column temperature was maintained at 30°C, with a flow rate of 1 mL/min. The mobile phase consisted of water containing 0.05% trifluoroacetic acid (TFA, Thermo Scientific, 85183) and methanol (MeOH, LC-MS grade, DAMAS-beta, 75851G). Detection was performed at 214 nm using a 40 min gradient elution program.

### LC-MS/MS analysis

LC-MS/MS analysis was performed using an Easy-nLC 1200 coupled with a Q Exactive mass spectrometer. Peptide separation was achieved using a 150 μm inner diameter × 170 mm C18-AQ column (1.9 μm particle size; packing material: Reprosil-Pur). For positive ion mode acquisition, the mobile phases consisted of buffer A (0.1% formic acid [FA], LC-MS grade, Thermo Scientific, 225583) and buffer B (0.1% FA in acetonitrile/water [80:20, v/v]; acetonitrile, LC-MS grade, Thermo Scientific, F22MAB201). Mass spectrometry data were acquired using a data-dependent acquisition (DDA) mode. MS1 spectra were collected at a resolution of 70,000 with a scan range of *m/z* 100-1500, and MS2 spectra were acquired at a resolution of 17,500 with a normalized collision energy (NCE) of 28. Raw data were searched against the UniProt database (Taxonomy ID: 5476) using a nonspecific search strategy. The mass tolerance parameters were set to 20 ppm for precursor ions and 0.02 Da for fragment ions.

### Statistical analysis

Statistical significance is indicated by asterisks in the figures (\**p* < 0.05, \*\**p* < 0.01, \*\*\**p* < 0.001, and \*\*\*\**p* < 0.0001). Statistical comparisons were performed using two-tailed unpaired Student’s *t*-tests, one-way ANOVA with Tukey’s correction, and log-rank (Mantel-Cox) test. All statistical analyses were performed using GraphPad Prism software (version 9). Data are presented as mean ± standard error of the mean (SEM) unless otherwise indicated.

