## Supplemental figures for "Pulmonary colonization of *Candida albicans* drives breast cancer lung metastasis via candidalysin"

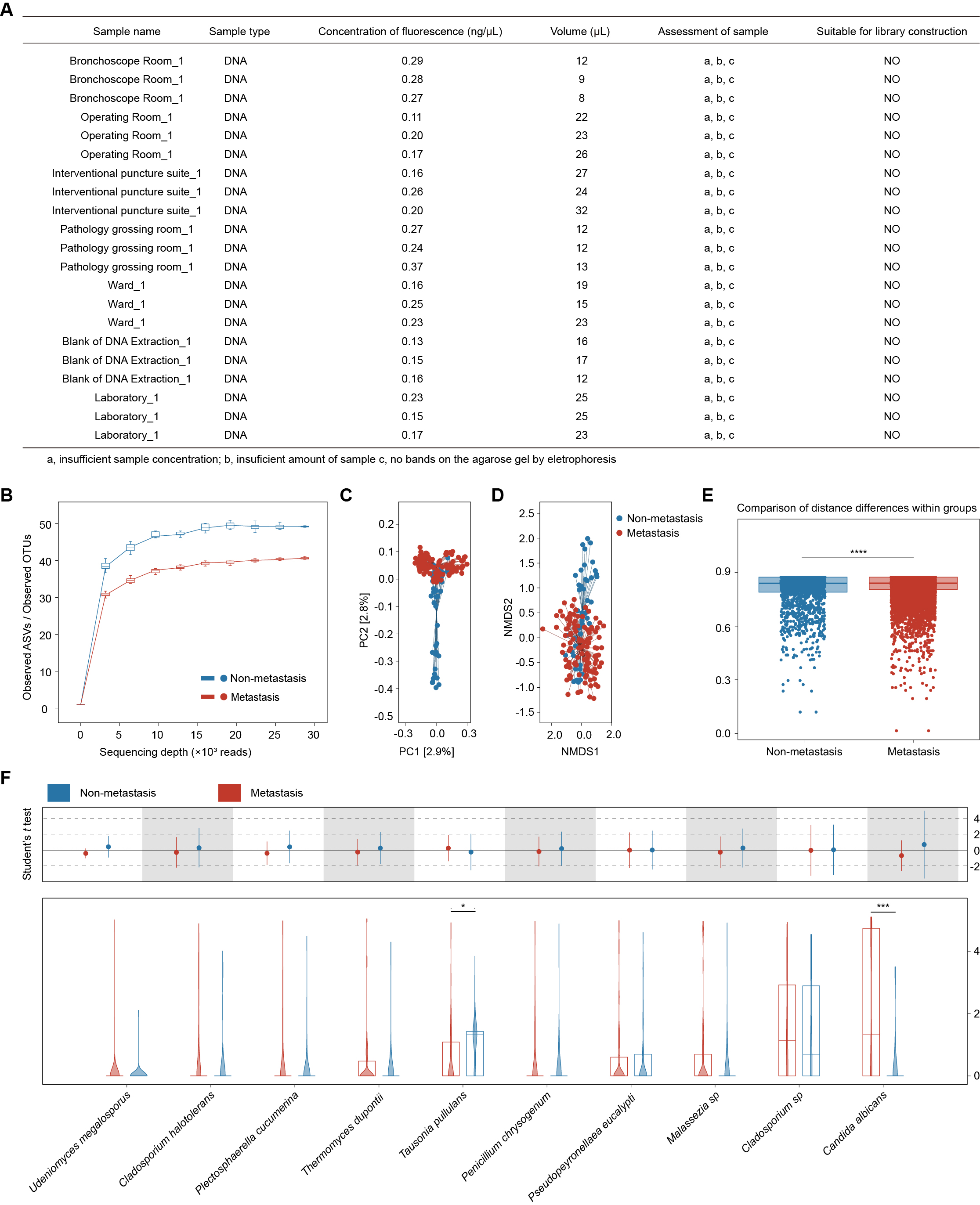
**Figure S1. Quality control of ITS sequencing and fungal microbiome diversity analysis, related to Figure 1**

(A) DNA concentrations of different environmental and blank controls and assessment of their suitability for library construction. Bronchoscope_N, blank control specimens collected from the bronchoscopy room; Surgery_N, blank control specimens collected from the operating room; Puncture_N, blank control specimens collected from the interventional puncture suite; Grossing_N, blank control specimens collected from the pathology grossing room; B_N, blank control specimens included during DNA extraction; Lab_N, blank control specimens collected from the laboratory; Ward_N, blank control specimens collected from the hospital ward.

(B) Rarefaction curves evaluating microbial alpha diversity and sequencing depth.

(C) Principal coordinates analysis (PCoA) plot showing differences in fungal community composition between the non-metastasis group (n = 45, blue) and the metastasis group (n = 112, red).

(D) Non-metric multidimensional scaling (NMDS) plot based on NMDS1 and NMDS2 axes, revealing spatial clustering and dispersion patterns of samples from the non-metastasis and metastasis groups in two-dimensional space.

(E) Quantitative comparison of within-group sample distances (within-group dissimilarity/beta dispersion) between the non-metastasis and metastasis groups.

(F) Relative abundance distributions and statistical comparisons of representative differentially abundant fungal species between the non-metastasis (blue) and metastasis (red) groups.

Data are represented as mean ± SEM. *p < 0.05; ***p < 0.001; ****p < 0.0001 compared using unpaired Student’s *t* test (two-tailed) (E and F).

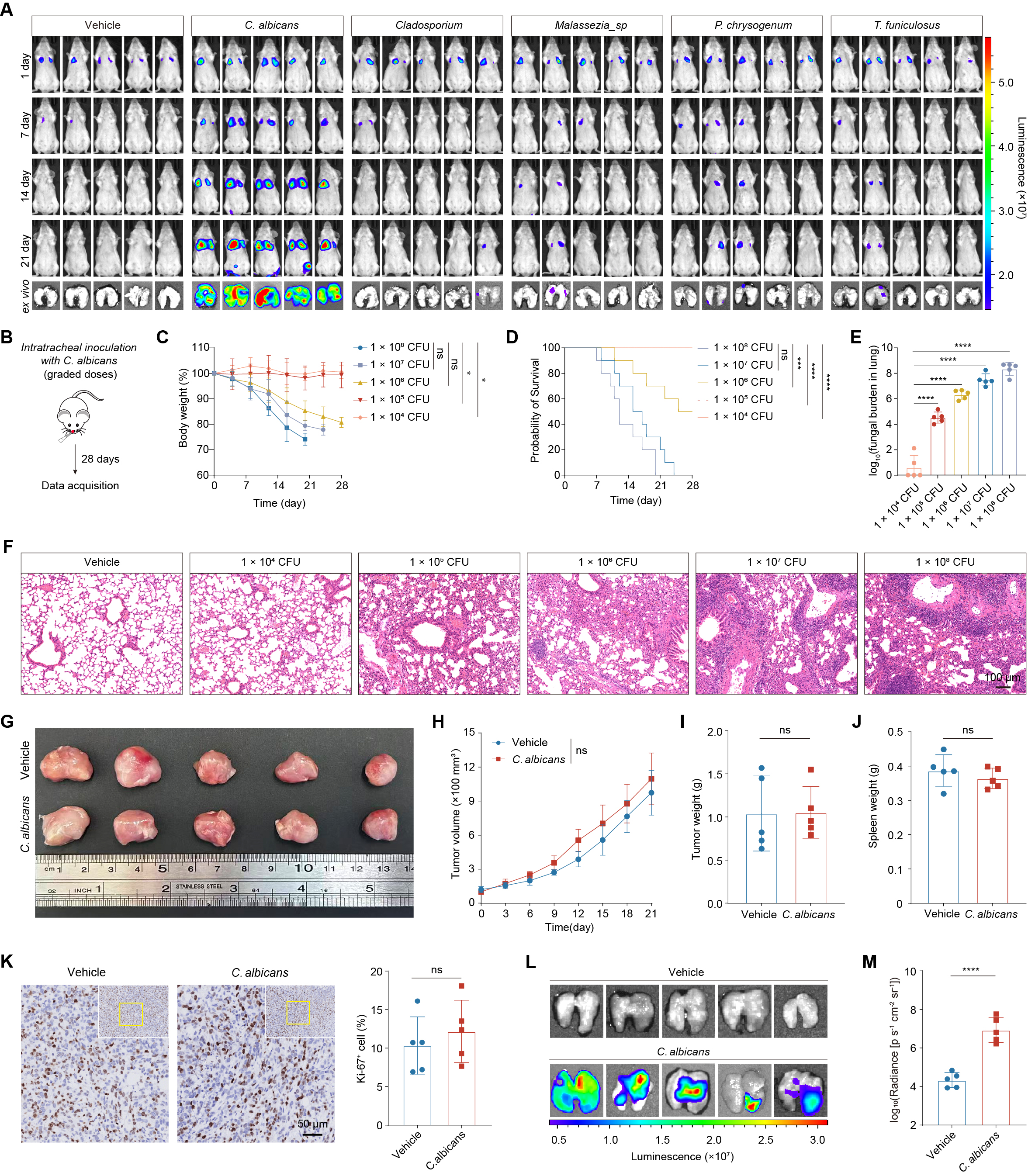

**Figure S2. Pulmonary *C. albicans* colonization promotes lung metastasis without significantly affecting primary tumor growth, related to Figure 1**

(A) Representative *in vivo* and *ex vivo* BLI images of pulmonary metastases following tail-vein injection of 4T1-Luc cells. Related to Figure 1F.

(B–F) Screening and optimization of the intratracheal *C. albicans* inoculation dose. (B) Schematic of the murine model of intratracheal *C. albicans* inoculation. (C) Body weight changes over 28 days in mice inoculated intratracheally with different doses of *C. albicans* (CFU/mouse). (D) Kaplan-Meier survival curves. (E) Quantitative analysis of pulmonary fungal burden in mice receiving different inoculation doses, expressed as log₁₀ (CFU). (F) Representative H&E-stained lung sections showing lung tissue damage and inflammatory cell infiltration induced by different doses of *C. albicans*. Scale bar, 100 μm.

(G–K) Effect of intratracheal *C. albicans* colonization on orthotopic primary tumor growth and spleen weight. (G) Images showing tumors collected from control and *C. albicans*-colonized mice. (H) Tumor growth curves. (I) Quantitative analysis of primary tumor weight at the endpoint. (J) Quantitative analysis of spleen weight at the endpoint. (K) Representative Ki-67 immunohistochemical staining and corresponding quantification. Scale bar, 50 μm.

(L–M) Representative *ex vivo* BLI images of pulmonary metastases (L) and longitudinal quantification of BLI signals (M).

Data are presented as mean ± SEM. ns, not significant; ****p < 0.0001 compared using unpaired Student’s *t* test (two-tailed) (H-K, and M), one-way ANOVA (C and E), and log rank (Mantel-Cox) test (D).

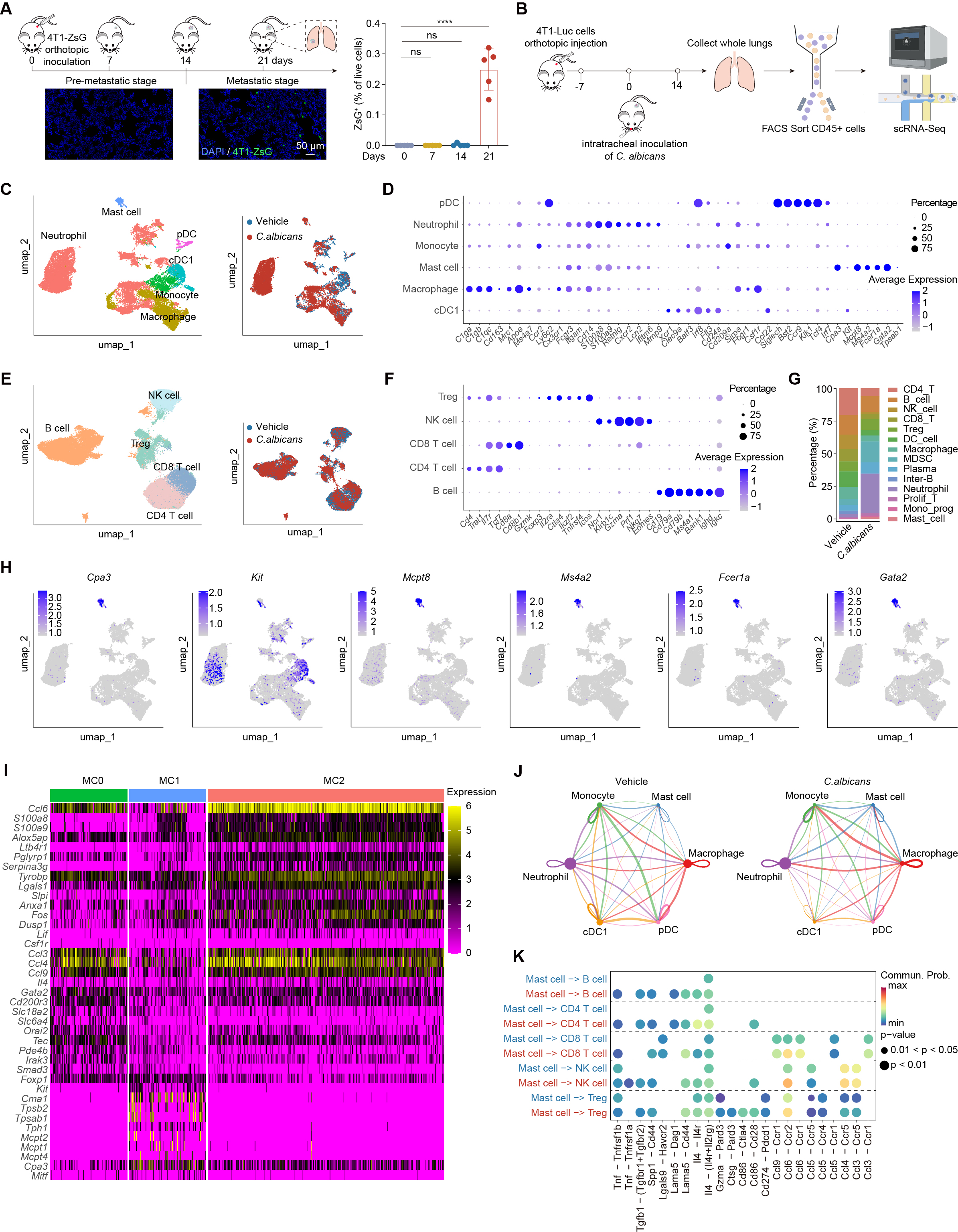

**Figure S3. Single-cell profiling of pulmonary immune subsets and mast cell subclustering in *C. albicans*-colonized lungs, related to Figure 2**

(A) Schematic of the pre-metastatic and metastatic stages in the orthotopic 4T1 tumor model (top left). Representative fluorescence micrographs of 4T1-ZsG cells in pulmonary sections from mice during pre-metastatic and metastatic stage (bottom left). Scale bar, 50 μm. (right) Flow cytometric quantification of ZsG^+^ tumor cell frequencies in the lungs at indicated time points, defining the pre-metastatic and metastatic phases.

(B) Schematic of pulmonary CD45^+^ cells isolation and scRNA-seq workflow in control and *C. albicans*-colonized mice.

(C) UMAP embedding of myeloid cells analyzed by scRNA-seq, showing cell type annotation and tissue origin.

(D) Dot plot showing the normalized average expression of canonical marker genes across individual myeloid cell subsets.

(E) UMAP embedding of lymphocyte analyzed by scRNA-seq, showing cell type annotation and tissue origin.

(F) Expression profiles of canonical marker genes across lymphoid lineages and related immune cell subsets.

(G) Bar plot showing the relative proportions of individual immune cell subsets.

(H) UMAP Visualization of mast cell marker gene expression.

(I) Heatmap showing marker gene expression levels in each mast cell cluster.

(J) Circular plots illustrate the interaction network of myeloid cells. Line thickness represents the overall signaling strength.

(K) Dot plot showing CellChat-inferred intercellular signaling from mast cell clustersto lymphocyte clusters (receivers).

Data are presented as mean ± SEM. ns, not significant; ****p < 0.0001 compared using one-way ANOVA (A).

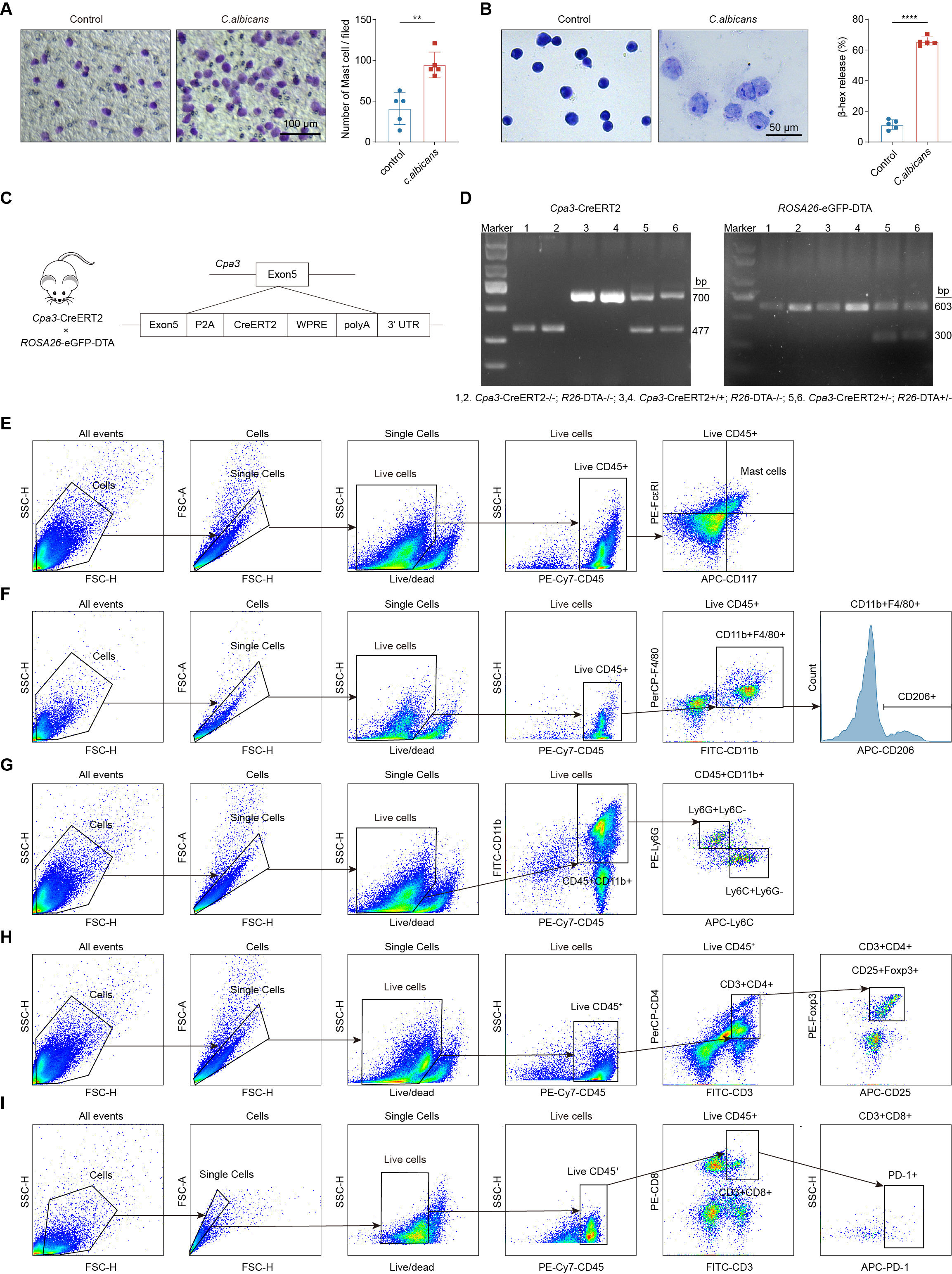

**Figure S4. Validation of *Cpa3*-DTA mice and flow cytometry gating strategies for pulmonary immune subsets, related to Figure 3**

(A) *C. albicans* promotes mast cell migration. Representative micrographs (left) and quantification (right) showing the number of mast cells migrating across the Transwell membrane in control and *C. albicans*-stimulated groups. Scale bar, 100 μm.

(B) Mast cell activation following *C. albicans* stimulation. Representative micrographs (left) and quantification of β-Hex release (right) in control and *C. albicans*-stimulated cells. Scale bar, 50 μm.

(C) Schematic illustrating the generation of conditional mast cell-depleted (*Cpa3*-DTA) mice.

(D) PCR genotyping on tail DNA of *Cpa3*-CreERT2⁻/⁻; *ROSA26*-DTA⁻/⁻ mice (1, 2), *Cpa3*-CreERT2⁺/⁺; *ROSA26*-DTA⁻/⁻ mice (3, 4), and *ROSA26*-CreERT2⁺/⁻; Cpa3-2A-CreERT2-DTA⁺/⁻ mice (5, 6) for *Cpa3*-CreERT2 (WT at 477 bp and *Cpa3*-CreERT2 at 700 bp) and *ROSA26*-DTA (WT at 603 bp and *ROSA26*-DTA at 300 bp).

(E–I) Gating strategies for flow cytometric analysis of major immune cell subsets in mouse lung tissues. (E) Mast cells, (F) M2 macrophages, (G) MDSCs, (H) Tregs, and (I) PD-1⁺ CD8⁺ T cells.

Data are presented as mean ± SEM. **p < 0.01; ****p < 0.0001 compared using unpaired Student’s *t* test (two-tailed) (A and B).

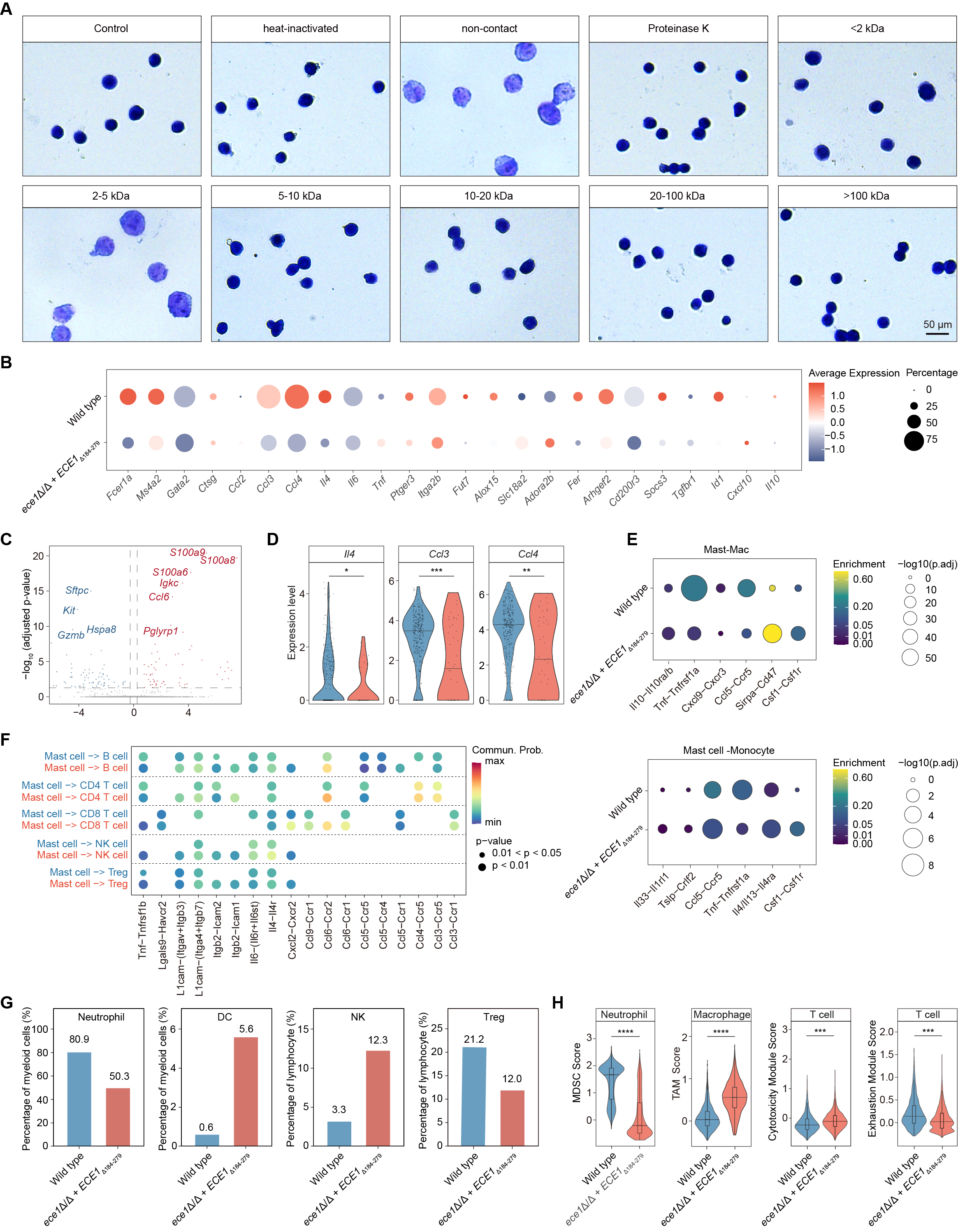

**Figure S5. Transcriptomic and cell-cell communication analyses of candidalysin-driven mast cell activation, related to Figure 4 and 5**

(A) Representative toluidine blue staining images showing the effects of different treatment conditions and molecular weight-fractionated supernatants on mast cell activation. Related to Figure 4B.

(B) Dot plot showing the expression patterns of key differentially expressed genes (DEGs) in mast cells between WT and *ece1*Δ/Δ + *ECE1*Δ_184-279_ *C. albicans*-colonized groups.

(C) Volcano plot showing differentially expressed genes in mast cells.

(D) Violin plots showing expression of key activation and inflammatory genes in mast cells.

(E) Ligand-receptor communication networks between mast cells and macrophages (top) or monocytes (bottom) predicted by CellChat.

(F) Dot plot showing CellChat-inferred intercellular signaling from mast cell clusters (senders) to lymphocyte clusters (receivers).

(G) Quantification of major pulmonary immune cell subsets in mice colonized with WT and *ece1*Δ/Δ + *ECE1*Δ_184-279_ *C. albicans*.

(H) Violin plots displaying functional gene module scores across immune cell subsets.

Data are presented as mean ± SEM. *p < 0.05; **p < 0.01; ***p < 0.001; ****p < 0.0001 compared using unpaired Student’s *t* test (two-tailed) (D and H).

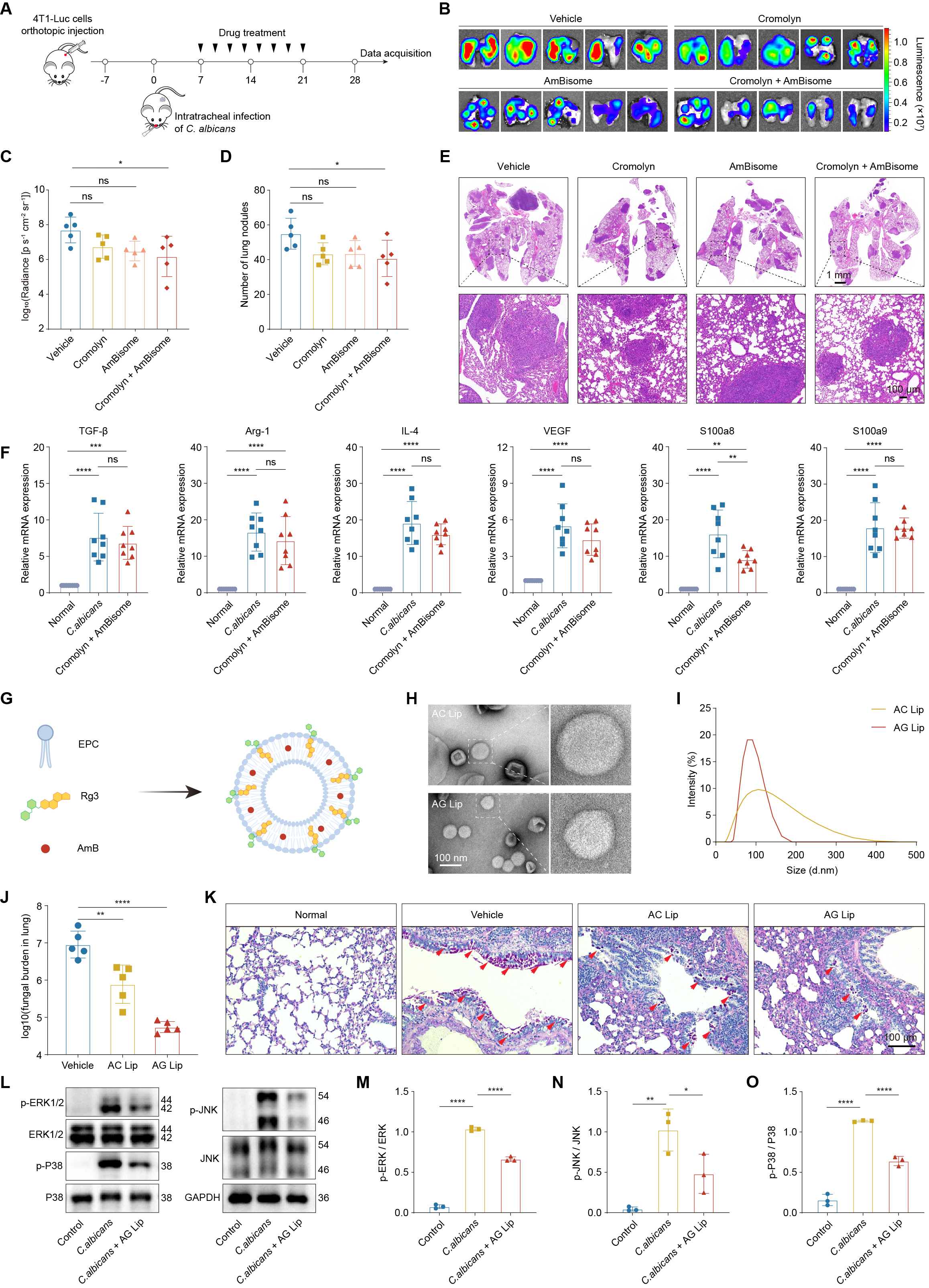

**Figure S6. Characterization of co-encapsulated liposomes and comparison with the free drug combination, related to Figure 7**

(A) Schematic of the mouse model and treatment regimen with conventional drug combinations.

(B–C) Representative *ex vivo* BLI images of pulmonary metastases (B) and longitudinal quantification of BLI signals (C).

(D–E) Quantification of pulmonary metastatic nodules at the endpoint (D) and representative H&E-stained images (E).

(F) qPCR analysis showing TGF-β, Arg-1, IL-4, VEGF, S100a8, and S100a9 expression after *C. albicans* colonization and Cromolyn + AmBisome treatment.

(G) Schematic of the design and preparation of ginsenoside Rg3 and amphotericin B co-encapsulated liposomes (AG Lip).

(H) Transmission electron microscope (TEM) images of AC Lip (top) and AG Lip (bottom).

(I) Hydrated particle size curves of AC Lip and AG Lip.

(J–K) Evaluation of fungal clearance and therapeutic efficacy after treatment with different amphotericin B formulations. (J) Quantification of pulmonary fungal burden. (K) PAS staining (arrows indicate fungal elements; scale bar, 100 μm).

(L) Immunoblots showing the protein expression levels of phosphorylated ERK1/2, JNK, and P38 in mast cells following *C. albicans* stimulation and AG Lip treatment. GAPDH was used as a loading control.

(M-O) Quantification of phosphorylated ERK1/2 (M), JNK (N), and P38 (O) relative to total proteins.

Data are presented as mean ± SEM. ns, not significant; *p < 0.05; **p < 0.01; ***p < 0.001; ****p < 0.0001 compared using one-way ANOVA (C, D, F, J, and M-O).
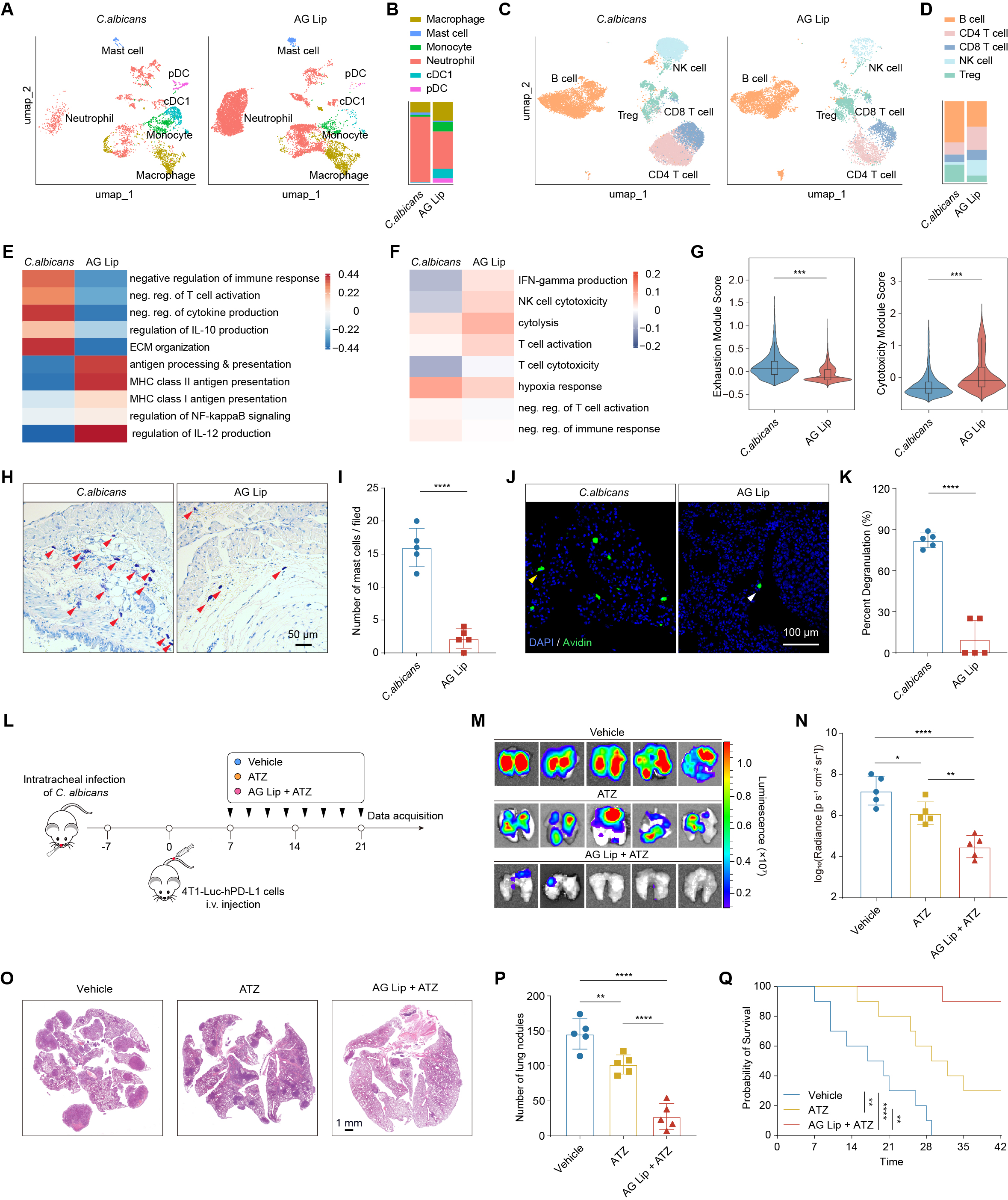
**Figure S7. Co-encapsulated liposomes synergize with PD-L1 blockade and reshape the pulmonary immune landscape, related to Figure 7**

(A) UMAP plots of pulmonary myeloid cells following *C. albicans* colonization and AG Lip treatment.

(B) Proportion of pulmonary myeloid cells.

(C) UMAP plots of pulmonary lymphocytes following *C. albicans* colonization and AG Lip treatment.

(D) Proportion of pulmonary lymphocytes.

(E–F) Heatmaps showing functional pathway analysis of myeloid (E) and lymphocyte (F) subsets.

(G) Violin plots showing the exhaustion (left) and cytotoxicity (right) module scores of T cells.

(H–I) Representative images of mast cell staining (H) and quantification of the number of mast cells per field (I). Red arrows indicate toluidine blue-positive mast cells. Scale bar, 50 μm.

(J–K) Representative immunofluorescence images of DAPI and Avidin staining (J) and quantification of percent degranulation (K). Scale bar, 100 μm. Yellow arrows indicate degranulated mast cells; white arrows indicate non-degranulated mast cells.

(L) Schematic of the lung metastasis model and treatment regimen.

(M–N) Representative *ex vivo* BLI images of pulmonary metastases (M) and quantification of luminescence (N).

(O–P) Representative H&E-stained images (O) and quantification of pulmonary metastatic nodules (P).

(Q) Kaplan-Meier survival curves.

Data are presented as mean ± SEM. *p < 0.05; **p < 0.01; ***p < 0.001; ****p < 0.0001 compared using unpaired Student’s *t* test (two-tailed) (G, I, and K), one-way ANOVA (N and P), and log rank (Mantel-Cox) test (Q).
